# A connexin 43 targeting peptide prevents blood vessel neointima formation

**DOI:** 10.1101/2025.09.22.677165

**Authors:** Mark C. Renton, Meghan W. Sedovy, Xinyan Leng, Farwah Iqbal, Melissa R. Leaf, Kailynn Roberts, Ramya C. Joshi, Arya Malek, Clare L. Dennison, Angela K Best, Paul D. Lampe, Joseph W. Baker, Mark Joseph, George S. Baillie, Brant E. Isakson, Scott R. Johnstone

## Abstract

The gap junction protein connexin 43 (Cx43) is associated with human pathological vascular smooth muscle cell (SMC) proliferation and neointima formation. We previously identified mitogen-activated protein kinase (MAPK) phosphorylation of Cx43 results in binding with the cell cycle protein cyclin E, facilitating neointima formation in mice. However, the specific nature of these interactions and their relevance to human disease have not been elucidated. Using an *ex vivo* human saphenous vein model of neointima formation, we identified increased MAPK-phosphorylated Cx43 and cyclin E in explant tissues. We used peptide arrays to define a cyclin E-Cx43 binding region and generated ‘CycliCx’, a stearate-linked Cx43 phospho-mimetic peptide. In human coronary artery SMC, CycliCx inhibits platelet-derived growth factor-BB (PDGF-β)-induced changes in Cx43 trafficking and interactions with cyclin E, and stimulation of proliferation. RNAseq analysis identified CycliCx significantly inhibits PDGF-β-induced proliferative pathways in SMC by limiting PDGF-β-induced early G1/S phase cell cycle progression transcripts. Finally, we show CycliCx limits neointima formation in mice *in vivo* and in *ex vivo* human saphenous vein explants. Our data provide strong evidence for selective targeting of Cx43 as a viable therapeutic strategy for preventing neointimal formation in humans.

## Introduction

Atherosclerosis in the coronary artery narrows the vessel lumen, restricting arterial blood flow, affecting more than 20.5 million Americans, making it the primary underlying cause of cardiovascular disease^1^. Surgical interventions, such as coronary artery bypass and percutaneous coronary interventions (angioplasty/stent placement), can be employed to revascularize and alleviate symptoms and risks of coronary artery disease. Of those who undergo surgical revascularization, 16.4% will experience a serious adverse event, including a heart attack or death^2^. This highlights the need for novel therapeutics to enhance vessel patency following revascularization surgeries.

The leading cause of long-term patency issues after coronary artery bypass grafting (CABG) or percutaneous coronary intervention is neointimal hyperplasia^3^, which occurs when vascular smooth muscle cells (SMC) in the medial layer of arteries/ vascular conduits dedifferentiate, take on a proliferative and migratory phenotype, and block the vessel intima^3^. Drug-eluting stents are often used to limit neointimal growth, yet these stents use broad antiproliferative agents, which delay vessel healing and necessitate the prolonged use of dual antiplatelet therapy, increasing the risk of clinically significant bleeding^3–5^. As such, identifying mechanisms of neointima development specific to SMC proliferation is essential in developing novel therapeutics.

Connexins form hexamers on the plasma membrane, which dock with connexin hexamers on adjacent cells, forming gap junctions, facilitating cell-cell communication^6^. Increasing evidence suggests connexins also regulate cell function in a gap junction-independent manner through protein-protein interactions^7–14^. In SMC, altered Cx43 expression has been associated with neointimal hyperplasia, but the specific role Cx43 plays remains unresolved, with conflicting studies showing both increased and decreased proliferative states in response to altered Cx43 expression^15–19^. Additionally, decreased expression and gap junction signaling are associated with both enhanced and delayed vascular disease progression^15,16,20–22^.

Nuclear magnetic resonance analysis of the cytoplasmic Cx43 C-terminus reveals two α-helical regions with a randomly coiled structure for the remaining length, forming a flexible protein primed for phosphorylation and protein-protein interactions^23^. In addition, Cx43 contains a long intracellular C-terminus with a high percentage of amino acids (a.a.) that can be post-translationally modified^10,24,25^. Phosphorylation state influences the conformation of the Cx43 C-terminus, altering the accessibility of other Cx43 phosphorylation sites and regulating the Cx43 protein-protein binding affinity^26^. Our previous study showed in response to platelet-derived growth factor-BB (PDGF-β), Cx43 is phosphorylated by mitogen-activated protein kinases (MAPK) at four of its C-terminal serines^24^. Phosphorylation by MAPK promotes Cx43 binding to the cyclin E protein, which binds with CDK2 to promote G1/S phase cell cycle transition^24,27^, and results in SMC proliferation in mice^24^. Our previous data suggested disrupting Cx43-MAPK activation does not compromise the viability of healthy SMC *in vivo,* making it a suitable target for therapeutic intervention^24^. In transgenic Cx43-MAPK phosphorylation-null mice, Cyclin E-Cx43 interactions were inhibited, and ligation-induced carotid artery neointima was reduced^24^. Despite this, the binding sites for Cx43 and cyclin E have not been elucidated.

In the present study, we aimed to define the importance of cyclin E-Cx43 interactions in human vascular tissues during neointima formation, and hypothesized that disrupting cyclin E-Cx43 protein-protein interactions could prevent neointima formation. We developed models of *ex vivo* human neointima formation using excess human saphenous veins intended for coronary artery bypass grafts and show alterations in cyclin E and Cx43 protein expression and phosphorylation during PDGF-β-induced SMC proliferation. We used peptide array techniques to define a 14 a.a., phospho-dependent cyclin E binding site on Cx43 and developed a stearate-linked linear peptide (‘CycliCx’) to disrupt cyclin E-Cx43 interactions. *In vitro* testing demonstrated CycliCx inhibits PDGF-β-induced changes in Cx43 localization and phosphorylation state, limits cyclin E-Cx43 interactions, and reduces human SMC proliferation. Transcriptomics data highlighted the CycliCx peptide disrupts PDGF-β-induced gene pathways regulating G1/S phase cell cycle progression in cultured human SMC. Importantly, our data show CycliCx is a potent inhibitor of neointima in mouse and human models. This study is the first to demonstrate specific binding domains for cyclin E-Cx43, show relevance in human disease, and identify a specific molecule targeted to SMC that limits neointima formation.

## Results

### A human saphenous vein model of neointima formation shows Cx43 MAPK-phosphorylation in new intima cells

Previous *ex vivo* models of human saphenous vein neointima developed by George et al.^28,29^ demonstrated high serum levels promote SMC proliferation and migration to the intimal surface of tissue explants, forming a “new intima” on the surface of the explants. We aimed to adapt this model to examine the effects of PDGF-β-induced proliferation using low serum media (**Figure 1A**). Human saphenous vein tissue explants underwent no treatment (NT; no PDGF-β) or 14-day treatment (PDGF-β, 50 ng/mL). Histological staining demonstrated an increase in cell layers at the surface of the explants in PDGF-β-treated tissue (**Figure 1B**, Supplemental Figure 1). The appearance of the cells is consistent with the new intima in previously published models for *ex vivo* HSV intimal growth^28,29^. Immunofluorescent staining demonstrated cells in the upper layers of the tissue are transgelin (TGLN) positive SMC, with significant increases in TGLN staining in PDGF-β-treated tissues (**Figure 1B**, Supplemental Figure 2). Staining also identified high levels of expression for total Cx43 and cyclin E within these cells, and antibodies specific to MAPK-phosphorylated sites on Cx43^24^ highlight PDGF-β promotes Cx43-MAPK phosphorylation in human tissues (**Figure 1B**, Supplemental Figures 3-5).

**Figure 1:**
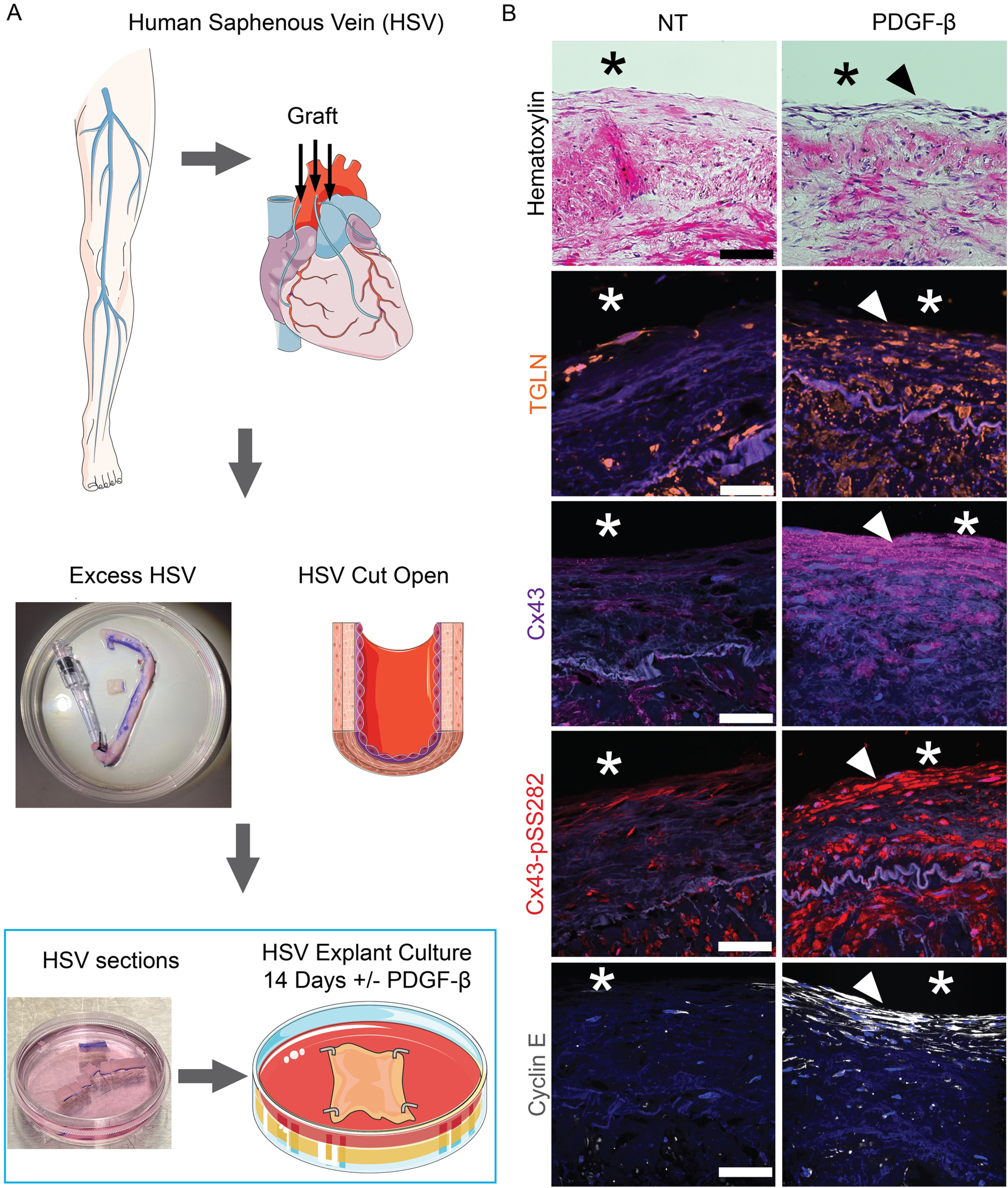
PDGF-β includes new intima in human saphenous veins in vitro, associated with an increase in Cx43 and proliferation markers

### Identification of a binding region between Cx43 and cyclin E and development of the CycliCx peptide

Our previously published data suggest phosphorylation at Cx43-MAPK C-terminal serines (S) regulates protein interactions between cyclin E and the Cx43 C-terminus^24^. However, specific sites of interaction are unknown, and structural modeling of Cx43 is limited. Therefore, we used a peptide array approach to define the critical binding sites of the Cx43 C-terminus required for binding with cyclin E. Peptide arrays of the full Cx43 C-terminus (a.a. 236-382) were designed with 25 a.a. peptide sequences arranged sequentially, with a five a.a. overlap between each sequence (**Figure 2A**). Arrays were arranged to test a requirement of the Cx43-MAPK serines located at S255/S262/S279/S282 (MK4) on Cx43, using either: non-mutated serines (Cx43^wt^, non-phosphorylated state), serine to alanine (A) mutations (Cx43^MK4A^, non-phosphorylatable state), serine to aspartate (D) mutations (Cx43^MK4D^, phosphorylated state mimic), or native phosphoserines (pS) (Cx43^MK4pS^, phosphorylated state). The full-length cyclin E protein was synthesized as we previously described^24^ (**Figure 2B**). Isolated cyclin E proteins were incubated with Cx43 peptide arrays, and binding was detected using antibodies targeting cyclin E (**Figure 2C-D**). Initial peptide arrays identified three potential binding regions in each of the Cx43^MK4D^ and Cx43^MK4pS^ sequences (**Figure 2C-D**). Given consensus between D/pS binding regions, we focused on a.a. 236-301, which were divided into three separate, but overlapping, 25 a.a. peptide sequences: Seq1 a.a. 235-260 (gray), Seq 2 a.a. 254-279 (green), Seq 3 a.a. 276-301 (black) (**Figure 2 E-F**). To test the requirement of single a.a., we prepared arrays using alanine substitutions along the length of each sequence, incorporating phosphorylation of Cx43-MAPK serines where they occurred. Our data indicate consistent binding when phosphorylation was present in Seq 1 and Seq 2, but no evidence of binding in Seq 3 (**Figure 2G**). Further, single a.a. substitutions did not significantly impact overall binding in Seq 1-2 (Supplemental Figure 6A).

**Figure 2:**
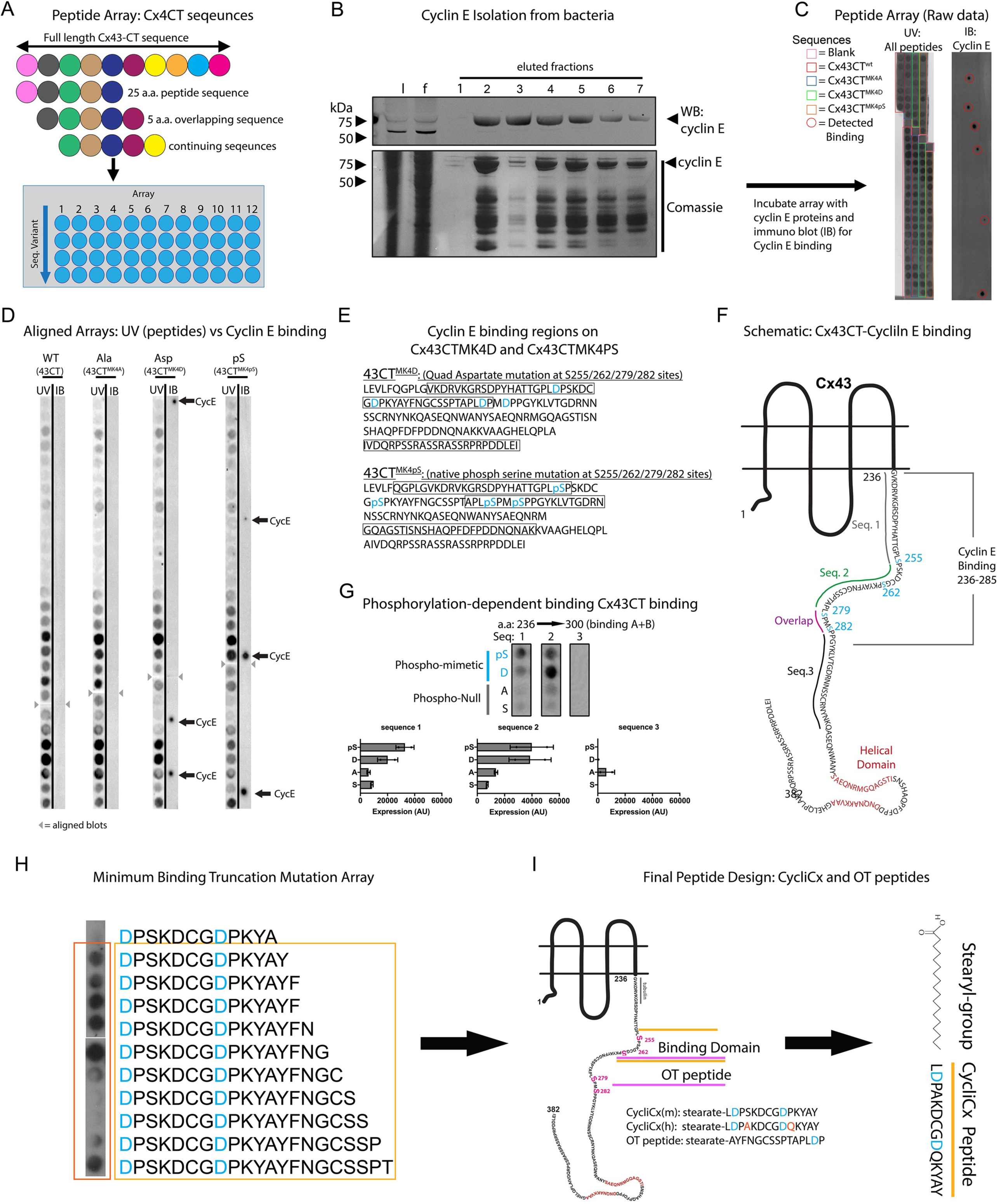
Design of the CycliCx peptide using peptide arrays demonstrates cyclin E binding regions on the phosphorylated Cx43CT

Using a truncation mutation peptide array approach, which reduces the sequence length by a single a.a. at a time, we tested regions of interest within the identified short region GVKDRVKGRSD (Peptide-1, 11 a.a.) and longer region LDPSKDCGDPKYAYFNGCSSPT (Peptide-2, 22 a.a.) that showed cyclin E binding (Supplemental Figure 6B-C). We synthesized peptides for these sequences, conjugated with an N-terminal stearyl group to aid in peptide cellular internalization. Peptides (30 μM) were incubated with cell cycle-stalled primary human coronary artery smooth muscle cells (CASMC) treated with PDGF-β (50 ng/mL) to promote proliferation. We used 5-ethynyl 2’-deoxyuridine (EDU) to label newly synthesized DNA and measured proliferation by flow cytometry. At 48 hours, PDGF-β increased proliferation in CASMC, which was reduced by Peptide 2 without notable cell loss (Supplemental Figure 6D). However, Peptide 1 treatment resulted in significant cell loss and was not tested further (Supplemental Figure 6D).

Peptide arrays indicated a minimum binding sequence within Peptide 2, between a.a. 254-268, but a loss of signal after removal of Tyrosine 267 (**Figure 2H**, Supplemental Figure 6C). Based on this, we created ‘CycliCx’, a 14 a.a. sequence, LDPSKDCGDPKYAY, which showed cyclin E-Cx43 binding. We also made a control off-target peptide (OT-peptide), a 15 a.a. sequence, AYFNGCSSPTAPLDP, which represents the immediately adjacent a.a. sequence unable to bind cyclin E. Both peptides were synthesized with an N-terminal stearate to promote cell entry, giving a final design of stearate-LDPSKDCGDPKYAY (CycliCx), and stearate-AYFNGCSSPTAPLDP (OT peptide), which was used as a control treatment throughout this study (**Fig 2H-I**). Because the CycliCx sequence was initially based on the mouse Cx43 sequence, we created a humanized CycliCx peptide, stearate-LDPAKDCGDQKYAY, for testing (**Fig 2I**). In testing, we saw no differences between mouse and human CycliCx peptides in their effects on cells (not shown). The human CycliCx peptide was used for all future experiments. The two peptides have approximately the same molecular weight and charge. The CycliCx peptide has a sequence predicted MW of 1571 kDa, (1853.22 kDa with stearate group) and a charge of -1, and the OT peptide has a predicted MW of 1586 kDa (1806.01 kDa with stearate group) and a charge of -1 (Supplemental Figure 6E-F).

### CycliCx peptide alters Cx43 cellular functions in human CASMC

Our previous data showed PDGF-β-induced ERK phosphorylation is a key factor in regulating cyclin E-Cx43 protein interactions in mouse primary SMC^24^. To assess ERK activation, human CASMC were treated with PDGF-β (50 ng/mL) to induce proliferation. Treatment with PDGF-β significantly upregulated ERK Threonine 202/Tyrosine 204 phosphorylation (pERK) in CASMC (**Figure 3A**, Supplemental Figure 7). Treatment of CASMC with either CycliCx or OT-peptide (30 μM each) did not significantly alter levels of PDGF-β-induced pERK in CASMC (**Figure 3A**, Supplemental Figure 7). In addition, total Cx43 expression was not affected by PDGF-β in CASMC. However, CycliCx altered Cx43 western blot protein banding (loss of upper phosphorylated bands), while OT-peptide had no effect (**Figure 3B**, Supplemental Figure 8). Based on previous reports of phosphorylation states present in each migrating band, we assessed Cx43 protein kinase-C (PKC) phosphorylation at Cx43-S368 (Cx43-pS368). Our data highlight CycliCx significantly increases phosphorylation at Cx43-S368 in CASMC (**Figure 3B**, Supplemental Figure 8). While most experiments in cells were performed at 16 hours, we found CycliCx produced alterations in Cx43-S368 phosphorylation as early as 3 hours (Supplemental Figure 9).

**Figure 3:**
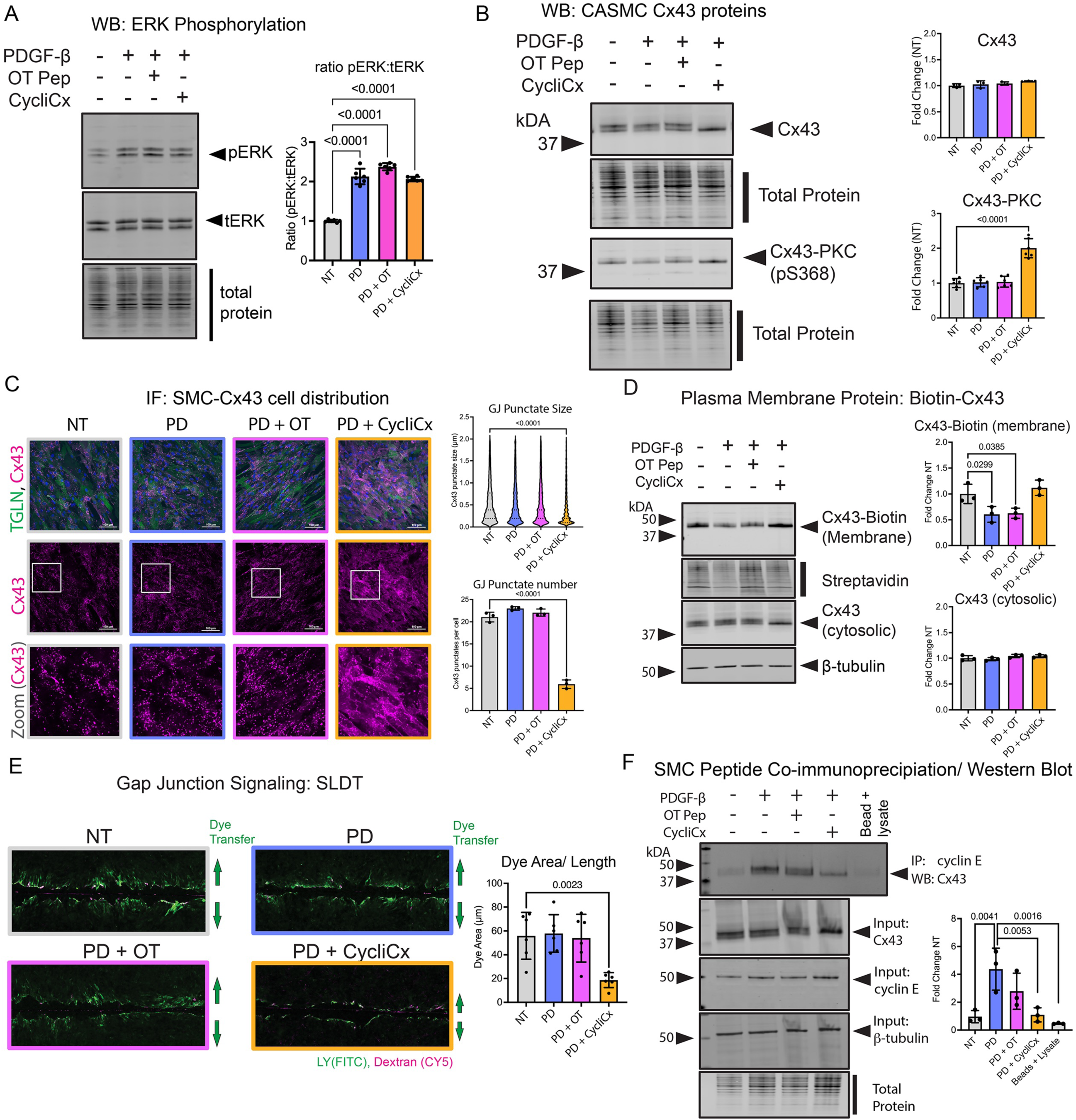
Design of the CycliCx peptide using peptide arrays demonstrates cyclin E binding

To assess Cx43 expression and localization, we performed immunofluorescence of cultured CASMC and identified punctate staining characteristic of Cx43, which was not significantly altered by treatment with PDGF-β or PDGF-β+OT peptide (**Figure 3C**, Supplemental Figure 10). In PDGF-β+CycliCx-treated cells, Cx43 GJ punctates were significantly reduced, and staining appeared more diffuse (**Figure 3C**, Supplemental Figure 10). Membrane biotinylation experiments demonstrated PDGF-β and PDGF-β+OT peptide treatments reduced Cx43 expression at the plasma membrane. CycliCx inhibited the PDGF-β-induced removal of Cx43 from the plasma membrane (**Figure 3D**, Supplemental Figure 11). To test whether CycliCx altered gap junction signaling, we performed scrape-loading dye-transfer assays using the gap junction-permeable lucifer yellow dye. We found CycliCx inhibited Cx43 dye transfer through gap junctions compared to controls (**Figure 3E**, Supplemental Figure 12). Finally, we tested if CycliCx could inhibit interactions between Cx43 and cyclin E. Co-immunoprecipitation assays demonstrated PDGF-β treatment increased cyclin E-Cx43 complex formation in human CASMC, which was significantly reduced by treatment with CycliCx (**Figure 3F**, Supplemental Figure 13).

### CycliCx peptide inhibits PDGF-β-induced proliferation by blocking G1/S phase cell cycle transition

To test if CycliCx alters the proliferation of SMC in a human model, we assessed PDGF-β-induced proliferation in CASMC by EDU flow cytometry. We found PDGF-β significantly upregulated CASMC proliferation, which was inhibited by treatment with CycliCx. (**Figure 4A-C**, Supplemental Figure 14). Bulk RNA sequencing of CASMC revealed PDGF-β+CycliCx altered the transcript expression of more genes than PDGF-β+OT peptide when compared with PDGF-β alone (**Figures 4D-E**). Pathway enrichment analysis using the Gene Ontology (GO) database demonstrated a significant upregulation of pathways involved in cell cycle progression and mitotic nuclear division in PDGF-β-treated compared to control CASMC, indicating CASMC differentiation into a pro-proliferative phenotype (**Figure 4F**). The addition of the CycliCx peptide resulted in significantly decreased enrichment of these same proliferative pathways when compared to PDGF-β treatment alone and to PDGF-β+OT peptide-treated CASMC (**Figure 4F**, Supplemental Figure 16).

**Figure 4:**
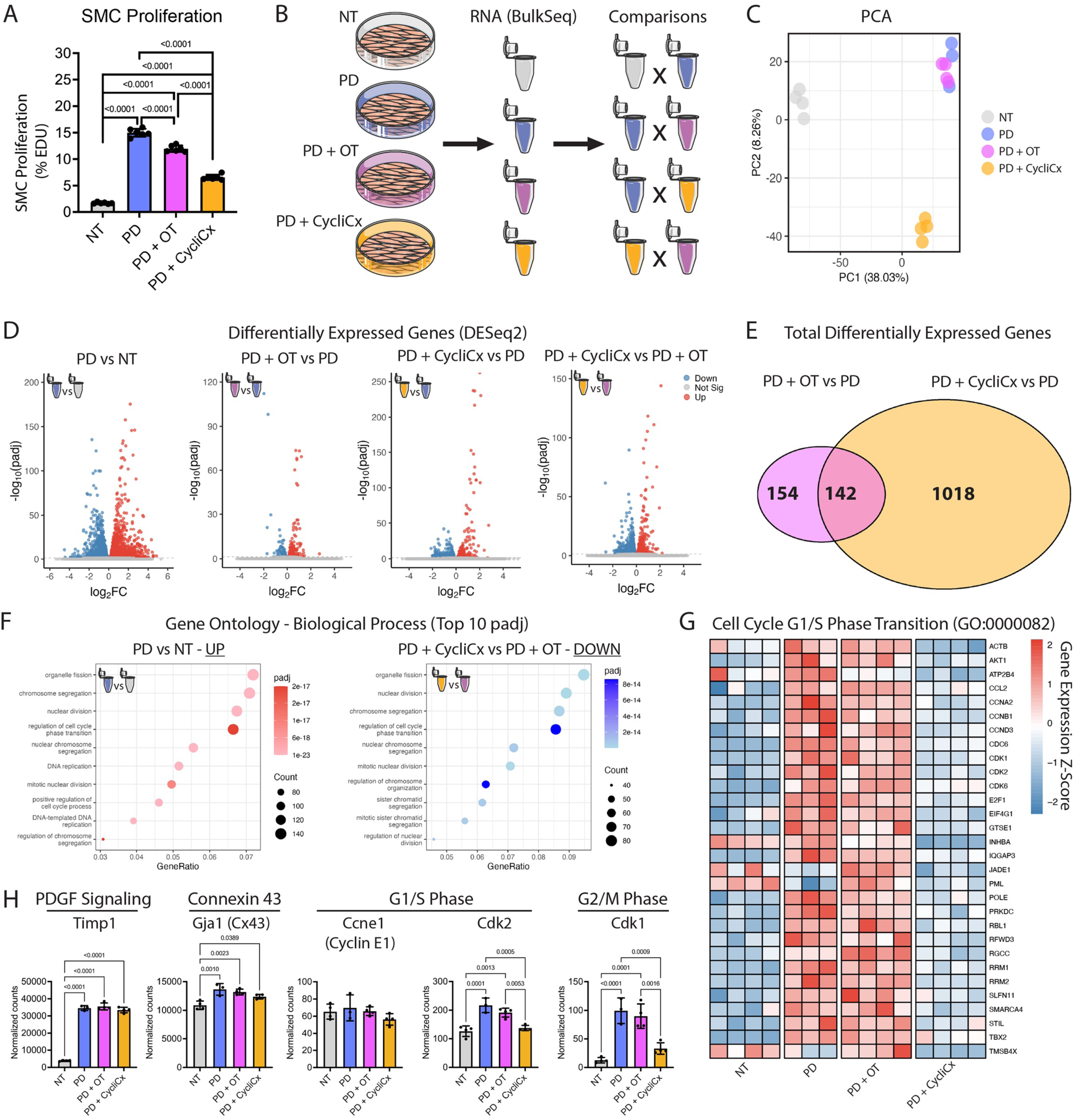
Transcriptions reveals that the CycliCx peptide arrests cell cycle progression at G1/S phase transition

**Figure 5:**
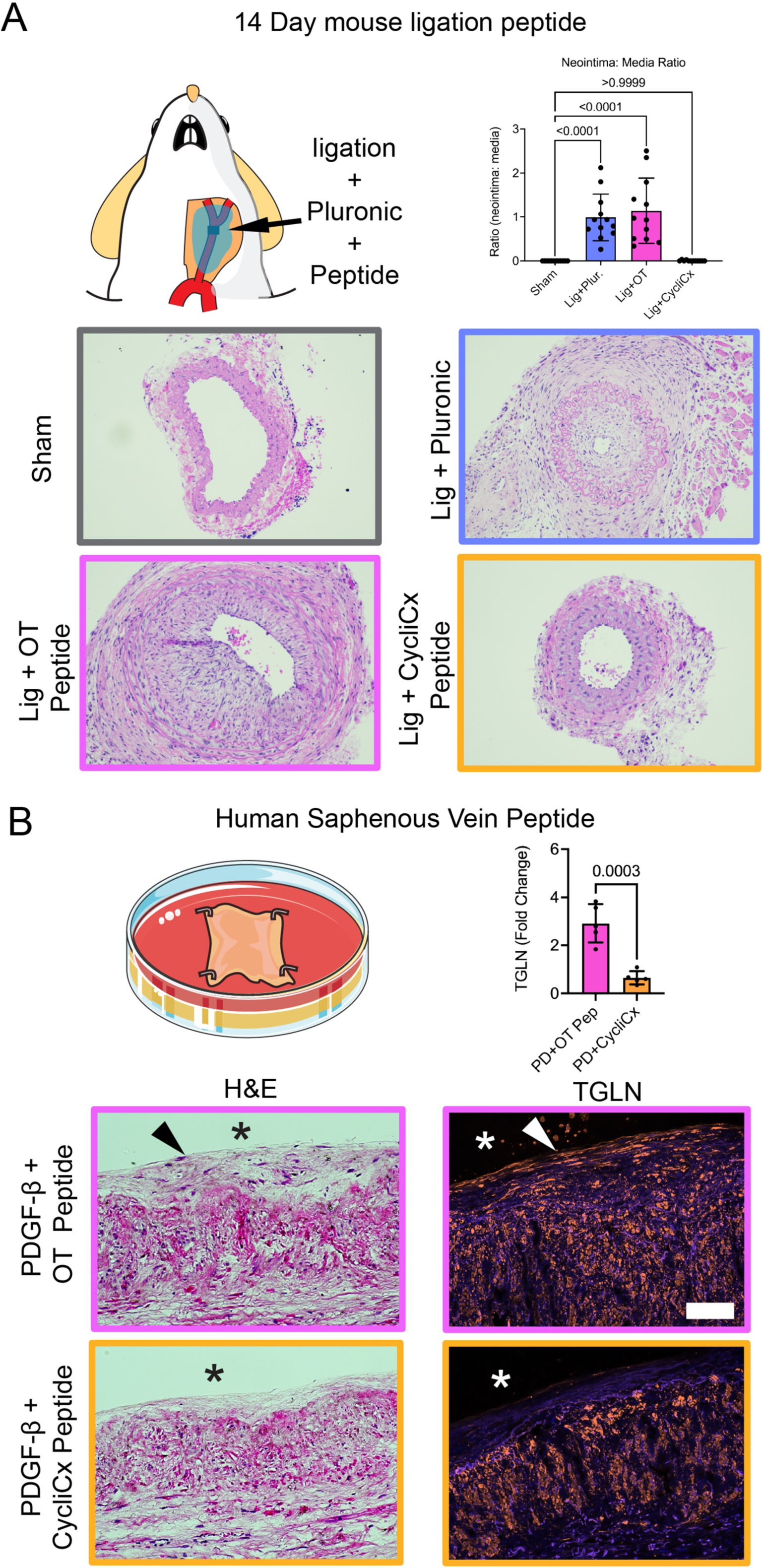
CycliCx peptide limits neointima formation in mouse and human blood vessels

Furthermore, CycliCx treatment significantly blunted the PDGF-β-induced transcript expression of large arrays of genes involved in ‘cell cycle G1/S phase transition’ (GO:0000082; **Figure 4G**), ‘cell cycle G2/M phase transition’ (GO:0044839), and ‘mitotic nuclear division’ (GO:0140014; Supplemental Figure 16). Importantly, PDGF-β+OT peptide treatment did not significantly blunt the PDGF-β-induced expression of these genes. We next interrogated the expression of key regulatory genes at each phase of the cell cycle (**Figure 4G**; Supplemental Figure 17) and found CycliCx treatment had no effect on the PDGF-β-induced expression of Ccnd1 (Cyclin D1) and Cdk4, which regulate early G1 progression, but partially blunted Cdk6 and Cdkn2c (p18) expression (Supplemental Figure 17A). Although CycliCx impairs binding of Cyclin E, it had no effect on Ccne1 (cyclin E1) gene expression (**Figure 4H**). However, CycliCx significantly blunted the PDGF-β-induced increase in the expression of G1/S phase transition promoters Cdk2 and E2f1 and partially reversed the PDGF-β-induced decrease in expression of G1/S phase transition inhibitors Tp53 (p53) and Cdkn1a (p21) (**Figure 4H**; Supplemental Figure 17B). Finally, we assessed late G2/M phase regulatory genes and found CycliCx inhibited the PDGF-β-induced increase in Ccna2 (Cyclin A2) and Cdk1, while OT peptide treatment had no effect (**Figure 4H**; Supplemental Figure 17C). Last, CycliCx did not alter the expression of Cx43 (**Figure 4H**), other canonical PDGF-β transcription targets such as Timp1 (**Figure 4H**), Timp3, and Il1b, or the downstream effector kinases Mapk3 (Erk1) and Mapk1 (Erk2; Supplemental Figure 18B). Last, smooth muscle contractile markers were decreased by treatment with PDGF-β, but treatments with CycliCx or OT peptide did not prevent this PDGF-β-induced transcriptional change (Supplemental Figure 19).

### CycliCx peptide disrupts neointima formation in mice and human blood vessels

To test CycliCx *in vivo*, we performed carotid ligation surgeries in mice and applied peptides mixed in Pluronic gel directly to the outside of carotids (**Figure 6A**). We found significant neointima formation in pluronic controls and OT peptide-treated mice compared with sham ligation mice (**Figure 6A**). Treatment with CycliCx peptide significantly inhibited ligation-induced neointima in mice and was not significantly different from sham control mice (**Figure 6A**, Supplemental Figure 20). We found no sex differences in mice in terms of neointima formation or the effect of the peptide, and data shown represent both male and female mice (Supplemental Figure 20). We then tested the effect of CycliCx in PDGF-β-induced neointima formation in *ex vivo* human saphenous veins. As shown in **Figure 1B** and Supplemental Figures 1-5, PDGF-β promotes the formation of SMC-derived new intima regions at the surface of tissue explants compared to NT. This is significantly increased when compared with no treatment (NT) at 14 days and is unaffected by OT-peptide treatments (**Figure 6B**, Supplemental Figures 1-5). In *ex vivo* human saphenous veins, CycliCx significantly reduces the presence of SMC in the intimal region of explants, suggesting inhibition of neointima formation (**Figure 6B**, Supplemental Figures 1-5).

## Discussion

Targeted therapies to address the mechanisms of neointima formation are needed to reduce the risk of stent/bypass failure. Our previously published data identified phosphorylation of Cx43 at its C-terminus MAPK residues regulates SMC proliferation and neointima formation in mouse models^24^. This control of SMC proliferation is dependent on interactions between the Cx43 C-terminus and cell cycle-regulating protein cyclin E^24^. In the present study, we defined for the first time the binding region of cyclin E on Cx43 and demonstrated MAPK phosphorylation sites within this binding region are required for direct cyclin E-Cx43 interactions. We developed the CycliCx peptide, which mimics a short 14 a.a phosphorylated region of the Cx43 C-terminus within the MAPK region, and was designed to inhibit cyclin E-Cx43 binding. Our data highlight CycliCx alters Cx43 localization, signaling, and trafficking while limiting cyclin E binding with Cx43. Using a transcriptomic approach, we found CycliCx significantly blunts PDGF-β-induced cell cycle progression, leading to reduced proliferation in cultured human CASMC. Finally, we showed CycliCx significantly alters neointima formation in models of disease in mouse and human tissues. This work establishes critical binding sites on Cx43 required for SMC proliferation and identifies Cx43 as a viable therapeutic target for the prevention of human neointima formation.

Current strategies for mitigating neointima formation after stenting procedures primarily include the use of non-cell-specific drugs such as the Rapamycin analogs, Everolimus and Zotarolimus. These drugs work by targeting mTOR signaling, a pathway essential for protein, nucleotide, and lipid synthesis, ultimately regulating cell growth and division^30^. Although mTOR inhibitors can stop the pathological proliferation and migration of SMC, which form neointima, they present a risk to other cell types in the vasculature, particularly endothelial cells that protect the vessel from platelet aggregation and ultimately thrombus formation^4^. Our recent investigation revealed significant endothelial damage caused by standard preservation techniques during coronary bypass surgery, and highlighted the need for endothelial healing^31^. Targeting cyclin E-Cx43 interactions disrupts a key signal required for pathological SMC growth without impacting cell viability or global mTOR signaling. As such, this is a promising strategy for targeting neointima formation/restenosis without impacting healthy cellular healing functions and adversely impacting patient health.

Emerging research reveals Cx43 proteins can act independent of forming cell-cell gap junction channels to control cellular functions^7–13^. We previously described Cx43 interacts with the cell cycle control protein cyclin E in mouse SMC *in vitro* and *in vivo*. Cyclin E tightly controls G1/S phase cell cycle transition, which is critical for pathological SMC proliferation^27,32^. Our data in human saphenous vein explants showed both Cx43 and cyclin E are expressed in the new intima layers after PDGF-β stimulation, suggesting a role for Cx43 in the SMC proliferation in these tissues. Given our previous evidence that blocking these interactions is important in mouse neointima formation, we set out to define the role of cyclinE-Cx43 interactions in human neointima and SMC proliferation.

This is the first study to comprehensively define the location of cyclin E-Cx43 binding. We identified two regions on the Cx43 C-terminus with consistent cyclin E binding activity and used these to generate Cx43 mimetic peptides. The first region, used to generate peptide-1, spans Cx43 a.a. 235-245, which does not encompass any reported Cx43-MAPK phosphorylation sites. This peptide is highly similar to the Cx43 mimetic peptide juxtamembrane 2 (JM2)^33^. The JM2 peptide sequence is designed to cover the binding region between Cx43 and microtubule subunit β-tubulin^33^. JM2 enhances the Cx43 β-tubulin interactions, reduces Cx43 trafficking to the plasma membrane, and reduces Cx43 hemichannel ATP release^33,34^. In our *in vitro* model of PDGF-β-induced CASMC proliferation, the overlapping peptide 1 resulted in cell loss, and further testing of this peptide was not performed, as our goal is to limit CASMC from entering a proliferative phenotype while maintaining vessel health. This is in line with previous work in 4T1 tumor cells, in which JM2 was found to induce ROS-mediated apoptosis by disrupting mitochondrial trafficking^35^. However, no JM2 cytotoxicity was reported in healthy cultured human microvascular endothelial cells^33^. Conflicting cellular responses to the JM2/peptide 1 sequences could arise from differences in cell types, with CASMC more susceptible to cell loss than the endothelium from this peptide sequence. Duration of treatment and dose may also be responsible for the differential effects reported.

The second cyclin E-Cx43 binding region, used to generate peptide 2 and the final CycliCx peptide, spans Cx43 a.a. 254-275, with phosphomimetic aspartate mutations at the known Cx43-MAPK phosphorylation sites, S255 and S262. Cyclin-dependent kinase 1 (CDK1) and ERK5 are implicated in the phosphorylation of Cx43 at one or both of the S255 and S262 sites^36–38^, which correlate with gap junction internalization and cell division^24,36–39^. To include an appropriate control condition when testing the CycliCx peptide, we developed the OT peptide based on the sequence immediately adjacent to CycliCx (Cx43 a.a. 266-280). The OT peptide contains the known Cx43-MAPK phosphorylation site Cx43 S279 (but not the S282 site), and in line with the CycliCx peptide, the OT peptide contains a phosphomimetic aspartate at its corresponding residue and an identical N-terminal stearate group. The OT peptide sequence was selected as a control due to its proximity to CycliCx and lack of cyclin E binding activity. The OT peptide is similar to the first 15 a.a. of the 18 a.a. TAT-Cx43266-283 peptide published by Gangoso et al., with the exception of the a.a. 279 serine to aspartate mutation^7,40^. TAT-Cx43266-283 is a Cx43 mimetic based on the cSRC binding domain. cSRC perpetuates cell proliferation in some cancers, and evidence suggests binding with Cx43 inactivates cSRC and limits cancer growth. It is thought TAT-Cx43266-283 binds cSRC directly, mimicking the effect of full-length Cx43 and limiting glioblastoma growth in culture and *in vivo*^7,40^. In the present study, the OT peptide did not inhibit SMC proliferation or neointima formation. Due to the similarity between TAT-Cx43266-283 and our OT peptide, it is possible our OT peptide is lacking critical a.a. for cSRC binding, the aspartate mutation within the OT peptide prevents cSRC binding, or cSRC does not play a significant role in injury-induced proliferation in SMC.

Proposed mechanisms of action for previously published Cx43 mimetic peptides include binding between the Cx43 mimetic and a physically interacting protein, or direct binding between the Cx43 mimetic and Cx43 itself^7,40,41^. Our peptide array data demonstrate sequences for CycliCx and the cyclin E protein can bind directly. Although we do not present direct evidence of CycliCx-Cx43 interactions, CycliCx induced a shift in Cx43 molecular weight attributable to an increase in Cx43-pS368, a known PKC phosphorylation site. Cx43-pS368 phosphorylation is associated with control over gap junction open/closed states and has been observed with other Cx43 mimetic peptides, a phenomenon we summarized in our previously published review^42^. For example, the Cx43 mimetic peptide αCT1, developed by the Gourdie lab to disrupt Cx43/ZO-1 interactions, binds directly to the Cx43 C-terminus, causing an increase in Cx43-PKC phosphorylation^41^. Mutating αCT1 to interact only with ZO-1 and not directly with Cx43 did not produce increases in Cx43-pS368, indicating direct Cx43 binding is required for αCT1-induced Cx43-pS368^41^. As CycliCx induces a similar phosphorylation shift, it is possible CycliCx is also capable of directly binding the Cx43 protein.

Cyclin E exerts control over the cell cycle via binding with its kinase CDK2. Although the complete mechanism of action of cyclin E-CDK2 is uncertain, it is known this complex translocates from the cytoplasm to the cell nucleus to phosphorylate the retinoblastoma protein and exert its effect on cell division^43–46^. In our previous studies, cyclin E-Cx43 complexes are initially present in the cell membrane but bind Cx43 and are removed after prolonged PDGF-β treatment, demonstrating Cx43 can accompany cyclin E-CDK2 during translocation^24^. In the present study, CycliCx treatment retained Cx43 at the cell membrane after PDGF-β treatment, likely attributable to the shift toward Cx43-pS368 phosphorylation, which is associated with stabilizing Cx43 at the membrane^47,48^. Therefore, it is possible Cx43 is required to chaperone cyclin E to the cell nucleus to progress the cell cycle and induce proliferation, and disrupting the cyclin E-Cx43 complex interaction and trafficking prevents this.

In line with this, our transcriptomic data suggest CycliCx treatment blunts the PDGF-β-induced increase in transcript expression of critical cell cycle promoter genes downstream of the cyclin E-CDK2 complex. Cyclin E-CDK2 primarily regulates the G1/S phase transition of the cell cycle^27^. Indeed, there was little change in the expression of genes involved in the regulation of the early G1 phase of cell cycle progression with CycliCx treatment, while CycliCx significantly blunted the PDGF-β-induced regulation of G1/S phase transition and subsequent G2/M phase transition and M phase genes. This suggests CycliCx targets cyclin E to arrest the cell cycle during the G1/S phase transition.

In addition to halting pathological SMC proliferation and neointima formation, it is important to examine potential off-target effects of the CycliCx peptide. We found many PDGF-β downstream transcription targets unrelated to cell cycle, such as tissue inhibitors of metalloproteases (TIMPs) and interleukins, were not affected by the CycliCx peptide. Our investigation also revealed a reduction in gap junction communication following CycliCx treatment. This may be attributable to the observed increase in Cx43-pS368, which is associated with gap junction closure^49,50^. Interestingly, PDGF-β also induced gene expression of Cx40, which was reduced after treatment with CycliCx. Together, these data demonstrate CycliCx treatment specifically inhibits PDGF-β-induced pathological SMC cell cycle progression without significant disruption to other cell signaling pathways.

A caveat to this study is the use of human veins over arteries in testing CycliCx treatment in human tissues. In modern clinical practice, the internal mammary or radial arteries are preferred for coronary bypass surgeries over the saphenous vein due to lower levels of neointima formation^51–53^. Even so, the saphenous vein is still used more often than any other vessel for coronary bypass when additional bypass conduits are required^54^. As such, testing CycliCx in venous tissue is highly relevant to surgical practice. Currently, we are unable to trace the half-life of the CycliCx peptide due to a lack of available antibodies to CycliCx and the mechanism of synthesizing the peptide. The CycliCx peptide is synthesized with an N-terminal stearate group, limiting our ability to add a reporter such as GFP to the C-terminus. Attempts to add 6xHis tags to the peptide resulted in insoluble peptides and were not viable for testing (not shown). However, our data show clear effects *in vivo* in mice at 14 days, and our *in vitro* studies show CycliCx effects appear around 3 hours and remain at >16 hours, suggesting a peptide with good stability, although this remains to be fully elucidated. Our data also shows CycliCx limits gap junction signaling; thus, we are unable to distinguish gap junction and connexin-mediated effects directly in this study. However, our previous study found that Cx43-gap junction signaling did not play a significant role in PDGF-β-induced SMC proliferation in mice^24^. The CycliCx peptide may therefore also act as a promising tool for the specific regulation of gap junction signaling, which is not currently available.

In conclusion, we show that targeting Cx43 and cyclin E interactions with the novel Cx43 mimetic peptide CycliCx limits SMC proliferation and neointima formation in mice and human tissue. The findings from this study highlight new pathways important in neointima formation, which could be targeted to improve patency in human revascularization strategies. While our study focuses on pathological proliferation of SMC in blood vessels, future research should test whether CycliCx could be utilized in other diseases of pathological proliferation for which Cx43 has been implicated in^55^.

## Methods

### Sex as a biological variable

Male and female mice were used to assess sex as a biological variable in animal studies. Our data do not show differences in cell pathology or neointima formation in male vs female mice, and the results report combined data for both male and female mice. The sex of the individual mice is indicated in Supplemental Figure 20.

### Key Resources

Reagents, materials, and antibodies are all listed in Supplemental Table 1

### *Ex vivo* human saphenous vein model of neointima formation

We adapted a previously described *ex vivo* model of human saphenous vein neointima formation developed by George et al.^28,29^. At the end of surgical procedures for coronary artery bypass graft, excess, normally discarded saphenous vein tissue was collected. De-identified specimens from Carilion Clinic cardiovascular patients were processed within 1 hr of the end of surgery. Tissues greater than 2 inches in length, free from visible valves and branch regions, were used in the study. Tissues were placed into a container of harvest media (20 mM HEPES-buffered RPMI-1640, 0.225 mg/mL papaverine hydrochloride, 5 μg/ml amphotericin B, 20 IU/mL sodium heparin). Samples were transferred to a wash media (20 mM HEPES Buffered RPMI-1640 with 2 mM L-Glut, 8 μg/mL gentamicin, 1 IU/mL penicillin, and 1 μg/mL streptomycin). Tissues were cut into four consecutive explant sections and pinned on 6-well plates containing Sylgard 184 elastomer (**Figure 1**). Explants were maintained in a low-serum media (bicarbonate buffered RPMI-1640, 2 mM L-Glut, 8 ug/mL gentamicin, 1 IU/mL penicillin, 1 μg/mL streptomycin, and 2% FBS). The four treatment groups corresponding to four sections were: 1) no treatment; 2) PDGF-β (50 ng/mL, Millipore #01-305); 3) PDGF-β (50 ng/mL) plus OT peptide (30 μM); and 4) PDGF-β (50 ng/mL) plus CycliCx peptide (30 μM). Media were changed each day for 14 days. At the end of the experiments, tissue explants were fixed *in situ* in 4% PFA at 4°C overnight. To preserve integrity, tissues were encapsulated in 3% agarose before tissue processing and paraffin embedding by the FBRI Histology Core.

### Cx43 peptide arrays for cyclin E binding

SPOT synthesis of peptides and overlay experiments were carried out as described^56^. Arrays were used as an n=1 in most cases. However, each spot represents unique, but overlapping sequences, which are replicated multiple times across multiple blots. Full-length cyclin E-GST protein was synthesized as previously described by us^24^. Cyclin E-GST proteins for arrays were isolated using anti-GST beads, spin concentrated (Amicon, 10K MWCO, Millipore #UFC901008), and buffer exchanged to PBS 125 mM NaCl, pH 6.5 on PD10 columns (Millipore #GE17-0851-01). Peptide arrays were initially bathed in 100% ethanol, equilibrated in PBS-Tween (PBST, 125 mM NaCl, 0.1% tween, pH 6.5) for 10 min, then blocked in PBST-5% block (PBST with 5% milk) at room temperature for 2 hr. Arrays were washed in PBST, then incubated with cyclin E proteins (2.86 mg/mL) in PBST-1% block (1% Milk). Binding on arrays was detected using cyclin E primary antibodies in PBST-1% block, followed by secondary detection using rabbit HRP (Thermo Fisher #31458) in PBS-T-1% block for 1 hr at room temp, then developed by chemiluminescent substrate (SuperSignal, Thermo Fisher # 34579) on x-ray film in a dark room. Developed films were scanned for image analysis (cyclin E, Odyssey, LiCor). Arrays were imaged using UV to visualize total peptides (Odyssey, LiCor). Sequences for the truncation mutation array can be found in Supplemental List 1.

### Peptides

Peptides used in the studies were designed based on the peptide array minimal binding sequences shown in **Figure 2**. Peptides were synthesized with an N-terminal stearate group to improve cellular uptake^57^. Peptide final sequences used in the study are CycliCx: stearate-LDPSKDCGDPKYAY and Off-Target (OT) peptide: stearate-AYFNGCSSPTAPLDP. Peptide properties are described in Supplemental Figure 6. Patent #WO2021021716: “Compositions and methods for inhibiting neointimal formation” covers the discovery and use of peptide (CycliCx) and control peptides. Peptides were commercially synthesized (ThermoFisher custom peptide synthesis services) to >80% purity (Supplemental Figure 6F). Peptides were reconstituted at room temperature in sterile PBS (for cell culture) or saline (for *in vivo*) to a concentration of 2 mM and stored at -80°C for use. Peptides used in cell culture and explant culture experiments were prepared to a final concentration of 30 μM in media. Peptides for *in vivo* applications in mice were prepared to a concentration of 200 μM in a Pluronic gel (35% final concentration), with 50 μL of the Pluronic gel applied around the ligated carotids.

### Animals

All mice were cared for under the provisions of the Virginia Tech Animal Care and Use Committee and followed the National Institute of Health guidelines for the care and use of laboratory animals. All studies were performed in C57BL6/J mice; strain number #000664, purchased from Jackson Labs. Mice were bred in-house and used between 12-14 weeks of age.

### Carotid ligations and carotid Pluronic gel applications in mice

A 50% Pluronic gel stock was made by adding 3 g of Pluronic powder to 3 mL of ice-cold sterile saline and mixing under cold conditions until dissolved. Pluronic gels were sterilized by autoclaving. For surgeries, Pluronic stocks (50%) were mixed with sterile saline or with peptides in sterile saline to achieve a final concentration of 35% Pluronic. Samples were maintained on ice to maintain a liquid form. Permanent 14-day carotid ligations were used to promote neointima in mice, as we previously described^24^. Mice were, anesthetized using isoflurane, placed on a surgically sterile heating pad with isoflurane, and the neck area cleared of hair and sterilized. Mice received analgesic (Ethiqa 0.5 mL) proximal to the surgical site. An incision was made in the neckline of the mice, carotid exposed, and a 6/0 suture was secured to the carotid immediately before the bifurcation. In sham control mice, a 6/0 suture was passed under the exposed carotid and then removed. In ligated and sham mice, carotids were elevated, and cold Pluronic gel (50 μL) was applied evenly around the carotid using a positive displacement pipette. Carotids were held in place for around 10 seconds until the Pluronic gel solidified (as we previously described)^58^. The surgical site was then closed, and the skin sutured, followed by post-operative recovery. Comparisons were made to surgical controls (sham) and OT peptide controls.

### Cell culture

Primary human CASMC were sourced from American Type Culture Collection (ATCC, #PCS-100-02), ThermoFisher (#C0175C) and Lonza (#CC-2583). Initial stocks of cells were thawed and maintained in vascular smooth muscle cell basal media (ThermoFisher #M231-500) containing smooth muscle growth serum (SMGS, ThermoFisher # S00725). Cells were maintained between 70-90% confluency in culture for up to 16 cell divisions. For experiments, 5×10^4^ CASMC per mL were plated in each well of a six-well plate. CASMC were serum starved to stall cell cycle by washing in PBS and then incubating with vascular smooth muscle cell basal media containing 2% FBS for 72 hr prior to treatments. Media were changed every 48 hours. Stalled CASMC were induced with PDGF-β (50 ng/mL) and harvested at 16 hr for western blot or 48 hr for flow cytometry of proliferation. For peptide treatments, cells were treated simultaneously with PDGF-β (50 ng/mL) and peptide (30 μM).

### Flow cytometry

For studies of proliferation, CASMC were treated with 5-ethynyl-2’deoxyuridine (EDU, Thermo Fisher, #C10425) at the same time as PDGF-β and peptide treatments. At the end of experiments, CASMC were trypsinized, centrifuged (700 rpm, 5 min), then resuspended in 0.5 mL PBS. Cells were fixed by adding 70% ethanol while vortexing. EDU detection was performed as per the manufacturer’s instructions. Flow cytometry data were collected using a BD Accuri C6 flow cytometer, and data analyzed using FlowJo software. Proliferation was calculated as a percentage of EDU-positive cells to total cells.

### Immunofluorescence

Tissues fixed in 4% PFA were embedded in paraffin and sectioned. Antigen retrieval was performed (Universal R, antigen retrieval solution, Electron Microscopy Solutions) on deparaffined slides containing tissue sections and tissues blocked in a solution of PBS, 0.1% NP-40 substitute, 1% BSA, where required, BSA was replace with 5% anti-goat serum. Primary antibody incubations were performed overnight at 4°C, samples washed, and secondary antibodies applied.

### Imaging and analysis

Histological imaging was completed on a Nikon Ci-L histology microscope using tile scan/ Z-stack functions. Data for the quantification of neointima was performed by a researcher blinded to sample identification. All immunofluorescent images were captured using Nikon NIE confocal microscope (A1, with resonance scanning). Single images, maximum intensity Z-stack projection, along with tile scanning, were used to visualize samples. All quantitative analysis was performed using NIS Elements Advanced Research software (Nikon). Examples of NIE image processing tool use are shown in Supplemental Figures 2 **and 12**. To analyze new intima in TGLN-stained tissues, a line (16pt) was drawn at the luminal edge of the tissues using NIE software using the signal intensity tool. Background signal was set for individual images based on tissue autofluorescence, and the area under the curve (AUC) for the TGLN signal was divided by the length of the tissue. 5-6 images were taken across each tissue, and the AUC was averaged for these values. Representative patient TGLN images used for analysis are shown in Supplemental Figure 2. For each HSV experiment, consecutive tissue samples are used, and a fold change to the individual patient NT sample TGLN signal was made to calculate differences.

### Scrape load dye transfer (SLDT) assays

Serum-starved CASMC were treated with PDGF-β and peptides for 16 hr. Cells were washed in PBS with calcium and magnesium, and scratch wounds were created using a #10 scalpel. Cells were then incubated with the gap junction permeable Lucifer Yellow (420 mw, final concentration 0.05%, Thermo #2790937) and gap junction impermeable Rhodamine Dextran-647 (10,000 mw, final concentration 100 μg/mL ThermoFisher #2860812) for 5 min. Cells were then washed in PBS three times, fixed in 4% PFA for 20 min at room temperature, washed in PBS, and imaged for dye transfer (Nikon A1 confocal). Analysis of dye transfer area for each condition was performed using NI Elements Advanced Research (Supplemental Figure 12). Normalized data (to length) was presented as Area/Length of dye transfer.

### Western blotting and membrane protein biotinylation and coimmunoprecipitation

Western blotting, cell surface biotinylation, and co-immunoprecipitation studies were performed as we previously described^24,59^. Proteins for western blotting and cell surface protein biotinylation were harvested in PBS lysis buffer containing 5 mM EDTA, 0.5% sodium deoxycholate, 0.5% NP-40 substitute, 1 mM sodium orthovanadate, 10 mM sodium fluoride protease inhibitor cocktail (1:100, Millipore #P8340), phosphatase inhibitor cocktail 2 (1:100, Millipore #P5726), phosphatase inhibitor cocktail 3 (1:100, Sigma #P0044). For co-immunoprecipitation, 10 mM n-ethylmethylamine (Millipore, #291145) was added to the lysis buffer. Protein samples were quantified by BCA assay before loading, and equal loading was confirmed using β-tubulin and total protein assay (Revert 700 Total Protein Stain, LiCor). Membranes were developed using LiCor secondary antibodies, and imaged on a LiCor Odyssey scanner. Expression analysis was performed using Image Studio (LiCor). Values were normalized to total protein stain or β-tubulin, and changes were calculated as a fold change compared to non-treated controls.

### Bulk RNA sequencing

Serum-starved CASMC were plated, stalled for 72 hr, and treated with PDGF-β and peptides overnight. Cells were washed 2x in ice-cold PBS and then lysed in 350 µl of RLT lysis buffer. RNA was harvested (RNeasy Mini Kit, Qiagen #74104) according to the manufacturer’s instructions, including the optional DNA removal step. Extracted RNA was quantified using a NanoDrop One spectrophotometer (Thermo Fisher, USA). Aliquots of RNA (1.5 µg) were then diluted to equal concentrations (75 ng/µl) in RNase-free water.

RNA integrity analysis, library preparation, sequencing, raw read count generation, and differential gene expression analysis were performed commercially by Novogene (Sacremento, CA, USA) using their Human mRNA Sequencing pipeline. Full parameters, protocols, and quality control reports are available on request, but are explained briefly below:

RNA integrity was assessed using the 5400 Fragment Analyzer System (Agilent Technologies, CA, USA). mRNA library preparation was performed with poly A enrichment and library quality was verified using Qubit assay for quantification, Agilent Bioanalyzer for size distribution, and qPCR for molarity using adaptor-specific primers. Sequencing was performed on the NovaSeq X Plus (Illumina, USA) platform with 150 bp paired-end reads and a minimum of 6 G raw data. Raw sequencing data in FASTQ format were cleaned using custom in-house (Novogene) Perl scripts. Reads with adapter contamination, reads containing more than 10% uncertain nucleotides, and reads in which more than 50% of bases had a quality (Q) score below 5 were removed. At this point, an outlying sample in the PDGF-β group was removed from all further analysis due to significant AT/GC content separation (Supplemental Figure 15). Reads were then aligned to the human genome (hg38) using ‘HISAT2’ (v2.0.05). Raw read counts were generated using ‘featureCounts’ (v1.5.0-p3). Fragments Per Kilobase of transcript per Million mapped reads (FPKM) was derived from raw data by Novogene and used to calculate Pearson’s correlation coefficient to assess intra group and between group variability, and hierarchical clustering. However, for all downstream analysis raw count normalization was performed using ‘DESeq2’ (v1.44.0) in the R statistical analysis program (v4.4.1). To exclude low count genes prior to ‘DESeq2’ normalization, only genes with at least 10 counts in one sample were kept. Gene symbols were extracted based on provided Ensembl ID using ‘clusterProfiler’ (v4.12.0) and ‘org.Hs.eg.db’ (v3.19.1). ‘DESeq2’ normalized counts were log_2_ transformed to ensure a normal distribution. Euclidean distances were then calculated and principal component analysis (PCA) was performed using ‘dist()’ and ‘prcomp()’, respectively, in base R.

Differential gene expression analysis was performed by Novogene for every possible combination of sample groups using ‘DESeq2’ (v.1.20.0) with Benjamini-Hochberg multiple comparison adjustment. An adjusted p-value (padj) < 0.05 and an |log_2_(FC)| > 0 was set as the threshold for differentially expressed genes (DEGs). Volcano plots displaying -log_10_(padj) and log_2_(FC) were generated in R.

Pathway enrichment analysis was performed on DEGs with a padj < 0.05 with ‘clusterProfiler’ in R using the Gene Ontology (GO) database. The list of CASMC detected genes after low gene count exclusion was used as the background gene set. The GO terms with a padj < 0.05 following Benjamini-Hochberg multiple comparison adjustment were considered statistically significantly enriched. Separate pathway enrichment analysis was performed for upregulated DEGs (log_2_(FC) > 0) and downregulated DEGs (log_2_(FC) < 0). Individual GO term heatmaps were generated by extracting ‘DESeq2’ normalized counts for all DEGs (when comparing PDGF-β+CycliCx to PDGF-β+OT peptide) in indicated statistically significantly enriched pathways and calculating a gene expression z-score for each sample. Data were then visualized using ‘pheatmap’ (v1.0.12).

To interrogate changes in expression of single genes, ‘DESeq2’ normalized counts were extracted, and the mean normalized counts were plotted in GraphPad Prism 10. A 1-way ANOVA followed by Tukey’s multiple comparison testing was performed to assess changes in gene expression between groups. Unless otherwise stated, RNA sequencing data were visualized using ‘ggplot2’ (v3.5.1). Differential gene analysis and GO pathway analysis data sets are shown in Supplemental Tables 1-2.

### Statistical analysis

Statistical analysis was performed using Prism version 10.4.2. Samples were assessed for normal distribution using a Shapiro-Wilk test, and outliers were identified using the Prism robust regression and outlier removal (ROUT) test (Q =1%). 1-way or 2-way multiple comparison ANOVA followed by Dunnett or Tukey’s post-test was used for comparisons between >2 groups. The t-test was used for comparisons of 2 treatment groups. Kruskal-Wallis and Mann-Whitney non-parametric tests were used for groups that did not display normal distribution. A minimum of n=3 was used for all statistical analyses, and n values for each experiment are stated in the figure legends. P values are shown for all statistical analyses, with a P value of less than 0.05 considered significant. Data represent mean ± SD.

### Approval for the use of human excess vascular tissues

The Institutional Review Board (IRB) at Carilion Roanoke Memorial Hospital reviewed and approved the use of de-identified human saphenous vein tissues normally discarded at the end of coronary artery bypass graft surgeries. The Carilion Clinic IRB determined the research does not meet the definition of human subject research as outlined in 45 CFR 46.102(d). Patient data, health data, and identifiable information were not included with specimens.

## Data availability

All data generated and used for analysis and original data sets are freely available for further analysis. Bulk RNA sequencing data have been deposited in the Gene Expression Omnibus (GEO) with accession number GSE312484

## Author contributions

MCR, MWS, MJ, GSB, BEI, and SRJ designed research studies; MCR, MWS, XL, FI, MRL, KR, RCJ, AM, CLD, AKB, MJ, GSB, and SRJ conducted experiments and acquired data; MCR, MWS, XL, KR, RJ, AM, CLD, GSB, BEI, and SRJ analyzed data; PDL provided reagents; MCR, MWS, BEI, and SRJ wrote the manuscript; All authors approved the final version of the manuscript. A coin flip decided the listed order of co-first authors MCR and MWS; both authors contributed equally.

## Supporting information

Reagent list

Peptide array sequence list

DEG table

GO terms

## Acknowledgments

We gratefully acknowledge the resources and services provided Ms. Ruth Macleod at the University of Glasgow for the production of peptide arrays. Servier medical art, licensed under CC BY 4.0 https://creativecommons.org/licenses/by/4.0/, was used to create figure schematics. Funding for this research includes AHA 19CDA34630036 (SJ), AHA 23PRE1010870 (MS), AHA and the DC Women’s Board 25POST1410066 (MR), NIH-F31HL170721 (MS), NIH-R215R21HL168614-02 (SJ), VT-Proof of Concept (SJ), and the Seale Innovation Award (SJ). Medical Research Council grant MR/y00364/1 (GSB). NIH HL120840 (BEI), NIH HL137112 (BEI).

