## Supplementary material for "A connexin 43 targeting peptide prevents blood vessel neointima formation": Figure Legends

**Figure 1: PDGF-β induces new intima in human saphenous veins in vitro, associated with an increase in Cx43 and proliferation markers**. Schematic of human saphenous vein tissue harvesting and processing. Excess saphenous veins, used for coronary artery bypass graft, were dissected open with the luminal side up, cut free from valves and sutured vessels, and sequential sections were plated for testing. Media containing 2% serum and no treatment (NT) was compared with samples treated with PDGF-β (50 ng/mL), which was changed daily (**A**). Tissues were sectioned and stained for H&E, Tgln (orange), Cx43 (magenta), Cx43pS279 (red), and cyclin E (white) (**B**). N=3-5 individual patient veins used for staining. * indicates luminal side of the vessel, arrow indicates regions of new intima that were detected in tissues, scale bar is 50 μM. *Original images used in **B** from **Supp. Figs. 1-5.***

**Figure 2: Design of the CycliCx peptide using peptide arrays demonstrates cyclin E binding regions on the phosphorylated Cx43CT**. Schematic showing the design for peptide arrays, using a sequence corresponding to the 180 a.a. Cx43-CT (**A**). Sequential 25 a.a. sequences were printed to nitrocellulose; consecutive sequences were overlapped by 5 a.a. (**A**). Eluted fractions from GST columns for cyclin E-GST proteins expressed in bacteria, expression shown by western blotting (anti-cyclin E, upper blot) and coomassie staining (lower gel, **B**). Peptide arrays with Cx43CT$^{WT}$ (red box) or MAPK phosphorylated Cx43-S255/262/279/282 (MK4) substitutions with either alanine (Cx43$^{MK4A}$, blue), aspartate (Cx43$^{MK4D}$, green), or native phosphor-serine (Cx43$^{MK4pS}$, orange) were bound by cyclin E proteins and detected with anti-cyclin E (**C**). Potential binding sites highlighted by red circles (**C**) and cyclin E arrays aligned with UV imaging of total peptide (**D**). Each spot represents a unique sequence, N=1 blot, **C-D**. Three potential binding sequence regions in Cx43$^{MK4D}$ and Cx43$^{MK4pS}$ sequences are underlined (**E**). Analysis of the first three 25 a.a. binding sequences in Cx43 CT spanning Cx43 a.a. 236-300 shows phospho-specific binding within Cx43 a.a. 236-285 (**F**), locations shown in schematic (**G**). Each spot represents a unique sequence, N=1 blot (**G**). Truncation mutation peptide array across Cx43 a.a. 255-272 used to define a minimum binding region (**H**). Each spot represents a unique sequence, N=1 blot (**H**). A schematic showing the location of the binding domain and selected adjacent region used as an Off-Target (OT) peptide (**I**). The final design of the CycliCx peptide incorporating an N-terminal stearyl group (**J**). *Original array blots used for **E, G, and H** from **Supp. Fig. 6***.

**Figure 3: CycliCx Peptide alters Cx43 expression and interactions with cyclin E**. CASMC received no treatment (NT), were treated with PDGF-β (PD, 50 ng/mL) alone, PD in combination with Off-Target (OT) peptide (30 μM) or PD with CycliCx (30 μM) for 16 hr. Western blots were performed for total-ERK (tERK), phosphorylated-ERK (pERK phosphor ERK Thr202/Tyr204), and ratio of pERK: pERK calculated N=6 blots) (**A**), for total Cx43 (N=3 blots) and PKC-phosphorylated Cx43 (Cx43-PKC, pS368, N=6 blots) (**B**), and difference in expression from not treated (NT) calculated. ROUT outlier test, Shapiro-Wilk normality test, one-way ANOVA, Dunnett's multiple comparison test to NT. Distribution, size, and frequency of Cx43 GJ punctates (magenta) in treated CASMC (TGLN, Green) were captured by confocal microscopy, and differences in gap junction punctates/ cell were measured using NIS-elements software (N=3 biological replicates, average 3 images per replicate) (**C**). ROUT outlier test, Shapiro-Wilk normality test, one-way ANOVA, Dunnett's multiple comparison test to NT. Cell plasma membrane protein biotinylation western blots for membrane-bound Cx43 (streptavidin eluted) compared to cytosolic (non-biotin bound) proteins (N=3 blots) (**D**). ROUT outlier test, Shapiro-Wilk normality test, one-way ANOVA, Dunnett's multiple comparison test to NT. Cell-to-cell communication through gap junctions was assessed by lucifer yellow dye transfer (green) in scrape-wounded monolayers of CASMC after treatment with PDGF (50 ng/mL) or PDGF (50 ng/mL) plus OT/ CycliCx peptide (30 μM). Scrape load dye transfer (SLDT) of lucifer yellow (green) permeability was used to measure gap junction dye transfer, area of dye normalized to length, with co-loading of high molecular weight dextran (magenta) used to determine non-gap junction dye area (N=6 biological replicates) (**E**). ROUT outlier test, Shapiro-Wilk normality test, one-way ANOVA, Dunnett's multiple comparison test to NT. Co-immunoprecipitation western blots were performed to assess Cx43-cyclin E binding in CASMC following peptide treatment (IP cyclin E, WB Cx43) (N=3 blots), with loading control western blot for Cx43 or cyclin E proteins (**F**). Sample loading for western blots was normalized to total protein or β-tubulin expression, and comparisons were made to PDGF-treated cells. ROUT outlier test, Shapiro-Wilk normality test, one-way ANOVA, Dunnett's multiple comparison test to PDGF. Data displayed as mean ± SD. *Original blot/images/data graphs used for **A** (**Supp. Fig. 7**), **B** (**Supp. Fig. 8**), **C** (**Supp. Fig. 10**), **D** (**Supp. Fig. 11**), **E** (**Supp. Fig. 12**), **F** (**Supp. Fig. 13**).*

**Figure 4: Transcriptomics reveals that the CycliCx peptide arrests cell cycle progression at G1/S phase transition**. 5-ethynyl-2'-deoxyuridine (EDU) proliferation assay in human coronary artery smooth muscle cells (CASMC) treated with no treatment (NT), PDGF-β (PD, 50 ng/mL), PD + Off-Target peptide (OT, 30 μM), or PD + CycliCx (30 μM) (**A**). Shapiro-Wilk normality test, one-way ANOVA, and Tukey's multiple comparison test across all groups. Schematic showing the design for bulk RNA sequencing and group combinations for analysis (**B**). Principal component analysis (PCA) of 'DESeq2' normalized and $\log_2$ transformed counts (**C**). Volcano plots of 'DESeq2' differentially expressed genes (DEGs) identified with $|\log_2(\text{Fold Change (FC)})| > 0$ and an adjusted p-value (padj) < 0.05 for indicated group comparisons (**D**). Venn diagram of the total number of identified DEGs unique to and shared by PD + OT and PD + CycliCx peptide treatment groups when compared to PD treatment alone (**E**). Top 10 pathways by padj from Gene Ontology (GO) Biological Process pathway analysis ranked by gene ratio (left panel shows top 10 upregulated pathways in PD vs NT and right panel shows top 10 downregulated pathways in PD + CycliCx vs PD + OT; **F**). Heatmap of between replicate 'DESeq2' normalized gene expression z-scores for PD + CycliCx vs PD + OT DEGs from the significantly enriched GO pathway GO:0000082: 'cell cycle G1/S phase transition (**G**). 'DESeq2' normalized gene counts in selected genes (**H**). N=3-4 per group. ROUT outlier test, Shapiro-Wilk normality test, one-way ANOVA, Tukey's multiple comparison test across all groups. Data displayed as mean ± SD.

**Figure 5: CycliCx peptide limits neointima formation in mouse and human blood vessels**

H&E staining of mouse carotids following 14 days of sham surgery (no ligation) or ligation with either pluronic gel, pluronic+OT peptide or pluronic+CycliCx pep (200 μM). Measure of media area, neointima area, and ratio of neointima to media. ROUT outlier test, Kruskal-Wallis multiple comparison test to NT, N=12 (9 male, 3 female mice per group). H&E staining of human saphenous veins treated with PDGF+OT pep or PDGF+CycliCx for 14 days. Tgln staining of human saphenous veins treated with PDGF+OT pep or PDGF+CycliCx for 14 days. Fold change was calculated compared to NT (***Supp. Fig. 2***, *includes comparisons to NT and PDGF-treated*). N=5 individual patient veins. ROUT outlier test, Shapiro-Wilk normality test, and unpaired t-test were used for PD+OT peptide: PDGF+CycliCx. Data displayed as mean ± SD. *Original images for **A** (**Supp. Fig. 20**), **B** (**Supp. Fig. 2**).*

**Supplemental Figure Legends:**

**Supplemental Figure 1: Human saphenous vein H&E staining comparison.** Representative images of H&E-stained human saphenous veins for no treatment (NT), PDGF-β (PD), PD + Off-Target (OT) peptide, and PDGF + CycliCx. * indicates luminal side of the vessel, arrow indicates regions of new intima that were detected in tissues, scale bar is 50 μM. N=5 individual patient (pt.) veins. Blue box highlights images used in **Figure 1** and magenta box indicates images used in **Figure 5**.

**Supplemental Figure 2: Human saphenous vein smooth muscle cell transgelin staining**. In **A,** representative images of anti-TGLN-stained human saphenous veins for no treatment (NT), PDGF-β (PD), PD + Off-Target (OT) peptide, and PDGF + CycliCx. * indicates luminal side of the vessel, arrow indicates regions of new intima that were detected in tissues, scale bar is 50 μM. In **B-C**, Measurement of the leading edge (16 pt) TGLN staining (area under curve), normalized to imaged vessel length and averaged N=5-6 measures per tissue, N=5 individual patient (pt.) veins. ROUT outlier test, Shapiro-Wilk normality test, unpaired t-test were used for NT:PDGF, PD + OT peptide: PDGF + CycliCx, and ROUT outlier test, Shapiro-Wilk normality test, one-way ANOVA, Dunnett's multiple comparison test of all values. Data displayed as mean ± SD. Blue box highlights images used in **Figure 1,** and magenta box indicates images used in **Figure 5**

**Supplemental Figure 3: Human saphenous vein Cx43 staining comparison.** Representative images of anti-Cx43 stained human saphenous veins for no treatment (NT), PDGF-β (PD), PD + Off-Target (OT) peptide, and PDGF + CycliCx. * indicates luminal side of the vessel, arrow indicates regions of new intima that were detected in tissues, scale bar is 50 μM. N=3 individual patient (pt.) veins, Blue box highlights images used in **Figure 1**

**Supplemental Figure 4: Human saphenous vein Cx43-pS279 staining comparison.** Representative images of anti-Cx43-pS279 stained human saphenous veins for no treatment (NT), PDGF-β (PD), PD + Off-Target (OT) peptide, and PDGF + CycliCx. * indicates luminal side of the vessel, arrow indicates regions of new intima that were detected in tissues, scale bar is 50 μM. N=3 individual patient (pt.) veins, Blue box highlights images used in **Figure 1**

**Supplemental Figure 5: Human saphenous vein cyclin E staining comparison.** Representative images of anti-cyclin E1-stained human saphenous veins for no treatment (NT), PDGF-β (PD), PD + Off-Target (OT) peptide, and PDGF + CycliCx. * indicates luminal side of the vessel, arrow indicates regions of new intima that were detected in tissues, scale bar is 50 μM. N=3 individual patient (pt.) veins, Blue box highlights images used in **Figure 1**

**Supplemental Figure 6: Identification of active cyclin E binding sites on Cx43.** In **A**, Single alanine amino acid substitutions for Cx43 sequences Seq1 Cx43 a.a. 235-260 (white), Seq 2 Cx43 a.a. 254-279 (green), Seq 3 Cx43 a.a. 276-301 (black). Full-length sequences contain either native phosphoserine (ps) or aspartate (D). *The first four spots in each sequence contains the sequence with either serines (S), alanines (A), pS, or D, and were used as representative images for* ***Figure 2***. In **A**, arrays were imaged with UV (total peptides, left), then probed with anti-cyclin E, and detected with a secondary antibody before development by chemiluminescence onto x-ray film (right). Each spot represents a unique sequence, N=1 blot. In **B**, truncation mutation arrays were performed on sequences of interest (see **supplemental list** for sequences), detected with anti-cyclin E antibodies (top), and imaged with UV and Coomassie (total peptides). Each spot represents a unique sequence, N=1 blot. In **C**, alignments show the minimum binding sites identified across the Cx43-MAPK binding region (top, Sequence 2) and at a distant site (bottom, Sequence 1). *The top left blot in* ***C*** *is represented in* ***Figure 2***. Flow cytometry testing of proliferation using stearate-linked peptides targeting Sequence 1 and Sequence 2 incubated with SMC (30 μM) in the presence of PDGF (50 ng/mL) and EDU for proliferation (N=1 per treatment **D**). Histograms (*top,* **D**) and EDU data table (*bottom*, **D**). Sequence 1 reduced cell numbers, indicating cell death. Final design of shortened peptide sequences based on binding characteristics seen by truncation arrays for CycliCx peptide (mouse = m and human = h), and Off-Target peptide (OT-peptide, non-binding), approximate charge, and molecular weight not including stearate group determined by Expasy ProtParam tool (https://web.expasy.org/protparam/) (**E**). Example mass spec of commercially synthesized CycliCx (Top) and OT-peptide (bottom) used in experiments (**F**).

**Supplemental Figure 7: Full blots for ERK measurements**. CASMC received no treatment (NT), were treated with PDGF-β (PD, 50 ng/mL) alone, PD in combination with Off-Target (OT) peptide (30 µM) or PD with CycliCx (30 µM) for 16 hr. Western blots were performed for total ERK (tERK, p42/p44), phosphorylated ERK (pERK phosphor- p42/p44 Thr202/Tyr204) and ratio of pERK:pERK calculated (N=6). Total protein blots highlight equal loading of samples. Sample loading was normalized to total protein expression, and differences in expression were measured against non-treated cells. ROUT outlier test, Shapiro-Wilk normality test, one-way ANOVA, Dunnett's multiple comparison test to NT. Data displayed as mean ± SD. *Magenta box indicates blots/ graphs used in* ***Figure 3A****.*

**Supplemental Figure 8: Full blots for Cx43 measurements:** CASMC received no treatment (NT), were treated with PDGF-β (PD, 50 ng/mL) alone, PD in combination with Off-Target (OT) peptide (30 µM) or PD with CycliCx (30 µM) for 16 hr. Western blots were performed for Total Cx43 (N=3), Cx43-PKC (Cx43-pS368) (N=6). Sample loading was normalized to total protein expression, and differences in expression were measured against NT cells. ROUT outlier test, Shapiro-Wilk normality test, one-way ANOVA, Dunnett's multiple comparison test to NT. Data displayed as mean ± SD. *Magenta box indicates blots/ graphs used in* ***Figure 3B***.

**Supplemental Figure 9: CycliCx alters Cx43 phosphorylation expression from 3 hours post-treatment.** CASMC were treated with CycliCx (30 µM) and harvested at baseline and every hour up until 5 hr. Expression of total Cx43 and Cx43-PKC (Cx43-pS368) was measured. Data show a trend towards an increase in Cx43-PKC from approximately 3 hr post-treatment (N=2). Data displayed as mean ± SD.

**Supplemental Figure 10: IF of Cx43 expression in SMC following PDGF and peptide treatments.** CASMC received no treatment (NT), were treated with PDGF-β (PD, 50 ng/mL) alone, PD in combination with Off-Target (OT) peptide (30 µM) or PD with CycliCx (30 µM) for 16 hr. An N=3 images were taken per replicate (replicate N=3), and results were averaged in the analysis. *Highlighted boxes indicate images used in* ***Figure 3C***.

**Supplemental Figure 11: Full blots for plasma membrane protein biotinylation in CASMC**. CASMC received no treatment (NT), were treated with PDGF-β (PD, 50 ng/mL) alone, PD in combination with Off-Target (OT) peptide (30 µM) or PD with CycliCx (30 µM) for 16 hr. Western blots for Cx43 show plasma membrane-bound proteins (streptavidin eluted, upper blots) compared to cytosolic (non-biotin bound, Lower) proteins (N=3). ROUT outlier test, Shapiro-Wilk normality test, one-way ANOVA, Dunnett's multiple comparison test to NT. Data displayed as mean ± SD. Orange boxes indicate blots used in ***Figure 3D***.

**Supplemental Figure 12: IF of scrape load dye transfer in SMC following PDGF-β and peptide treatments.** CASMC received no treatment (NT), were treated with PDGF-β (PD, 50 ng/mL) alone, PD in combination with Off-Target (OT) peptide (30 µM) or PD with CycliCx (30 µM) for 16 hr. Monolayers of CASMC were scraped using a scalpel, followed by the addition of lucifer yellow (green) and Dextran (magenta). For each treatment group (NT, PDGF-β, PDGF-β + OT peptide, and PDGF-β + CycliCx) N=6 replicates were used (**A**). NIE image processing software was used to define the dye transfer area (**B**) and calculate the area of lucifer yellow gap junction dye spread minus the area of dextran used to detect scrape-injured cells (**C**). Both total dye area and dye area per scrape length were graphed (**D**). ROUT outlier test, Shapiro-Wilk normality test, one-way ANOVA, Dunnett's multiple comparison test to NT. Data displayed as mean ± SD. *Highlighted boxes indicate images used in* ***Figure 3E***

**Supplemental Figure 13: Full Co-immunoprecipitation western blots for Cx43 and cyclin E.** CASMC received no treatment (NT), were treated with PDGF-β (PD, 50 ng/mL) alone, PD in combination with Off-Target (OT) peptide (30 µM) or PD with CycliCx (30 µM) for 16 hr. Blots show cyclin E co-immunoprecipitation with Cx43 detected by western blot N=3). Control loading blots for Cx43, Cx43-GJA1-20K, cyclin E, β- tubulin, total protein. N=3, ROUT outlier test, Shapiro-Wilk normality test, one-way ANOVA, Dunnett's multiple comparison test to PDGF. Data displayed as mean ± SD. *Orange boxes indicate blots used in **Figure 3F***.

**Supplemental Figure 14: SMC EDU proliferation flow cytometry histogram data.** Flow cytometry histogram replicates for CASMC 5-ethynyl-2'deoxyuridine (EDU) fluorescence gating for cells that received no treatment (NT), were treated with PDGF-β (PD, 50 ng/mL) alone, PD in combination with Off-Target (OT) peptide (30 μM) or PD with CycliCx (30 μM) for 16 hr.. N=6 replicates, Shapiro-Wilk normality test, one-way ANOVA, Tukey's multiple comparison test to all samples.

**Supplemental Figure 15: Bulk RNA sequencing quality control, correlation, and clustering.** G/C and A/T content distribution along sequence reads for individual replicates in the PDGF-β (PD) group used as rationale for removal of PD_2 replicate from bulk RNA sequencing analysis. (**A**). Pearson's correlation coefficient matrix of FPKM normalized gene counts before (top panel) and after (bottom panel) removal of the outlying PD replicate (**B**). Heatmap of between replicate FPKM normalized gene expression z-scores with hierarchical clustering (**C**). N=3-4 per group. Con = no treatment control group; Pep4 = Off-Target (OT) control peptide; Pep5 = CycliCx peptide.

**Supplemental Figure 16: CycliCx peptide blunts mitotic gene pathways.** Heatmaps of between replicate 'DESeq2' normalized gene expression z-scores for PDGF-β (PD) + CycliCx vs PD + Off Target (OT) Peptide differentially expressed genes (DEGs) from the significantly enriched GO pathways GO:0044839: 'cell cycle G2/M phase transition (**A**), and GO:0140014 'mitotic nuclear division' (**B**). N=3-4 per group. NT = No Treatment control group.

**Supplemental Figure 17: CycliCx peptide alters the expression of critical cell cycle regulatory genes.** 'DESeq2' normalized gene counts from CASMC treated with no treatment (NT), PDGF-β (PD, 50 ng/mL), PD + Off-Target peptide (OT, 30 μM), or PD + CycliCx (30 μM) in selected genes that regulate early G1 phase progression (**A**), G1/S phase transition (**B**), and G2/M phase transition (**C**). Separated based on CycliCx regulation. N=3-4 per group. ROUT outlier test, Shapiro-Wilk normality test, one-way ANOVA, Tukey's multiple comparison test across all groups. Data displayed as mean ± SD.

**Supplemental Figure 18: Gene expression of gap junction proteins and PDGF regulators and transcription targets.** 'DESeq2' normalized gene counts from CASMC treated with no treatment (NT), PDGF-β (PD, 50 ng/mL), PD + Off-Target peptide (OT, 30 μM), or PD + CycliCx (30 μM) in selected gap junction genes (**A**), G1/S phase transition (**B**), and in PDGF regulatory kinases (Erk1/2) and non-cell cycle PDGF transcription targets (**C**). N=3-4 per group. ROUT outlier test, Shapiro-Wilk normality test, one-way ANOVA, Tukey's multiple comparison test across all groups. Data displayed as mean ± SD.

**Supplemental Figure 19: CycliCx peptide blunts the expression of smooth muscle contractile genes.** 'DESeq2' normalized gene counts from CASMC treated with no treatment (NT), PDGF-β (PD, 50 ng/mL), PD + Off-Target (OT, 30 μM), or PD + CycliCx (30 μM) in selected vascular smooth muscle cell contractile genes. N=3-4 per group. ROUT outlier test, Shapiro-Wilk normality test, one-way ANOVA, Tukey's multiple comparison test across all groups. Data displayed as mean ± SD.

**Supplemental Figure 20: Mouse ligation neointima measurements.** H&E staining of mouse carotid arteries following 14 days of sham surgery (no ligation) or ligation with either pluronic gel, pluronic + Off-Target (OT) peptide or pluronic + CycliCx peptide. Measure of media area, neointima area and ration of neointima to media. Highlighted images are used as a representative in **Figure 5**. Kruskal-Wallis multiple comparison test, N=12 mice (9 male, 3 female). Data displayed as mean ± SD. * indicates luminal side of the vessel, arrow indicates regions of new intima that were detected in tissues, scale bar is 100 µM. *Highlighted boxes indicate images used in **Figure 5A***
