## Supplemental Figures for "A connexin 43 targeting peptide prevents blood vessel neointima formation"

Supplemental Figure 1: HSV culture H&E

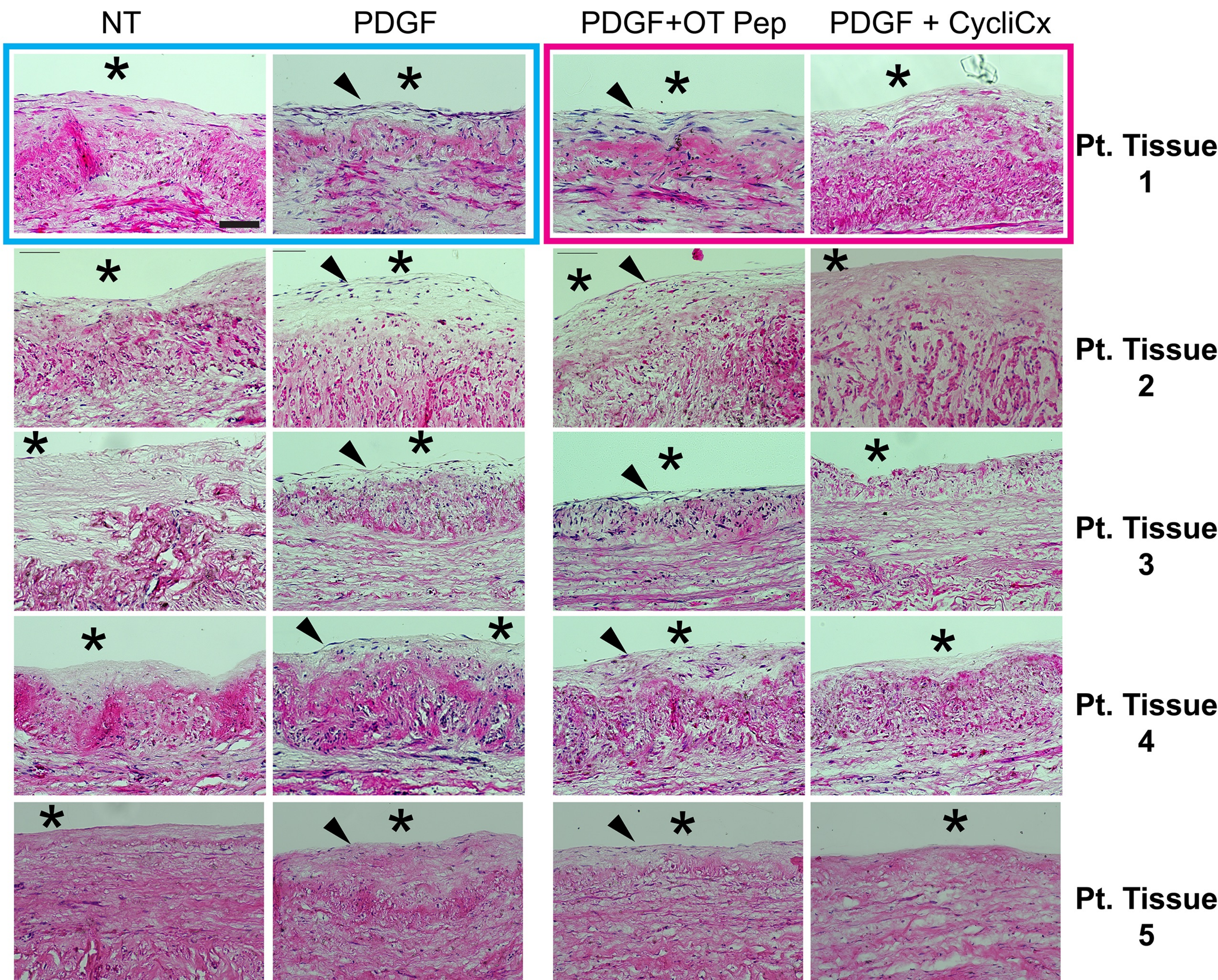

Supplemental Figure 2: HSV culture TGLN

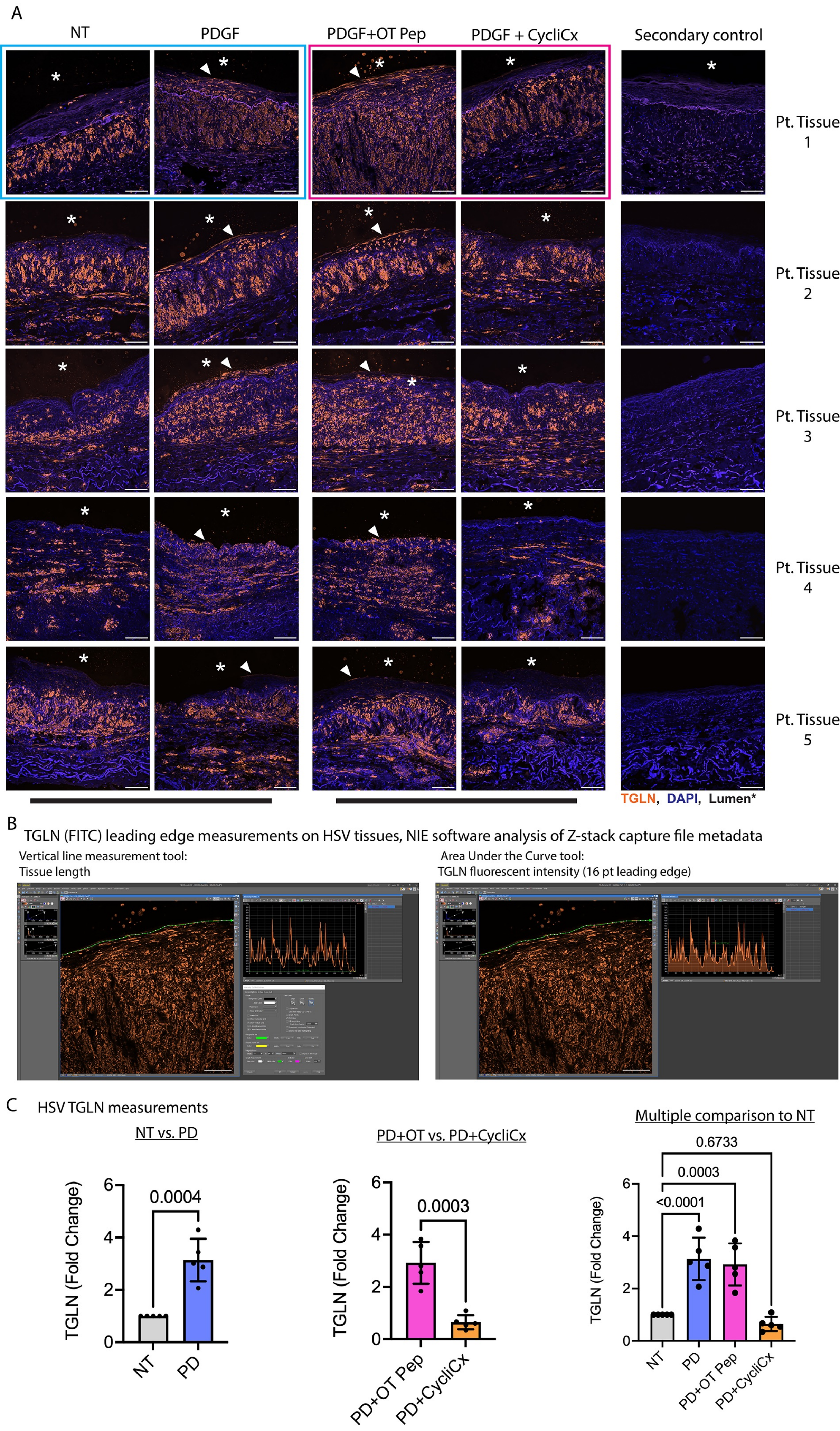

Supplemental Figure 3: HSV Culture Cx43

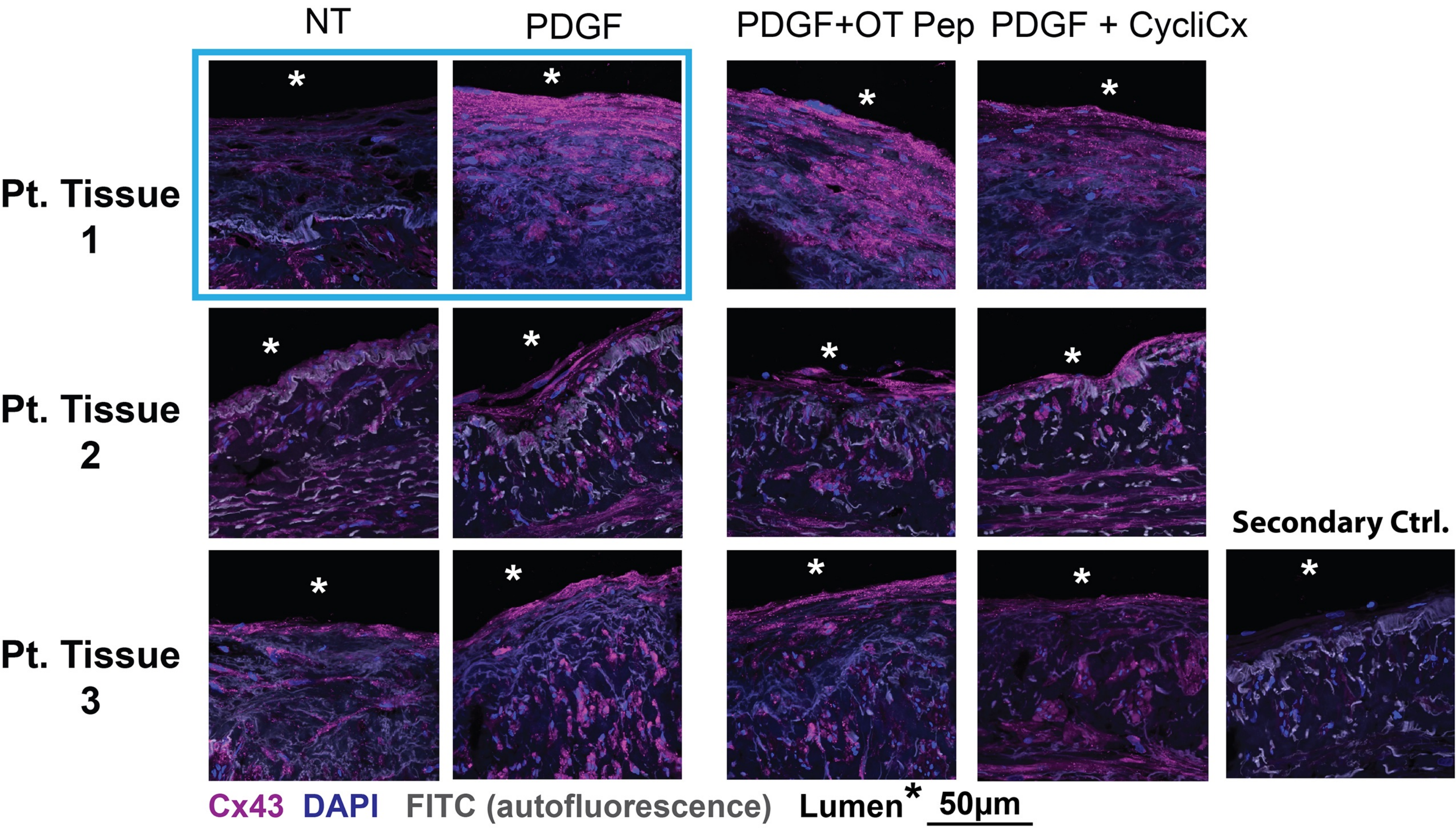

Supplemental Figure 4: HSV Culture Cx43-pS279

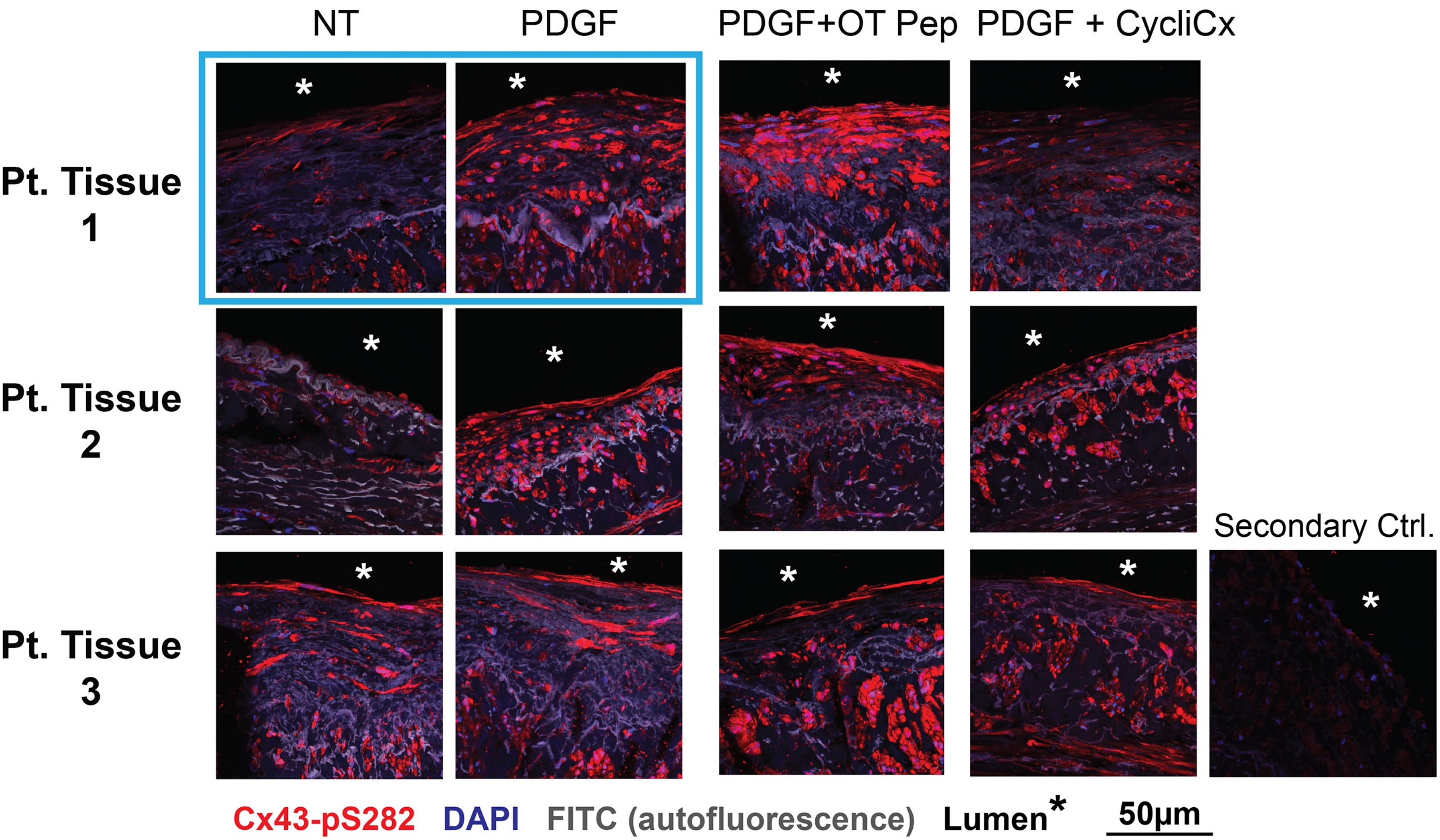

Supplemental Figure 5: HSV culture Cyclin E

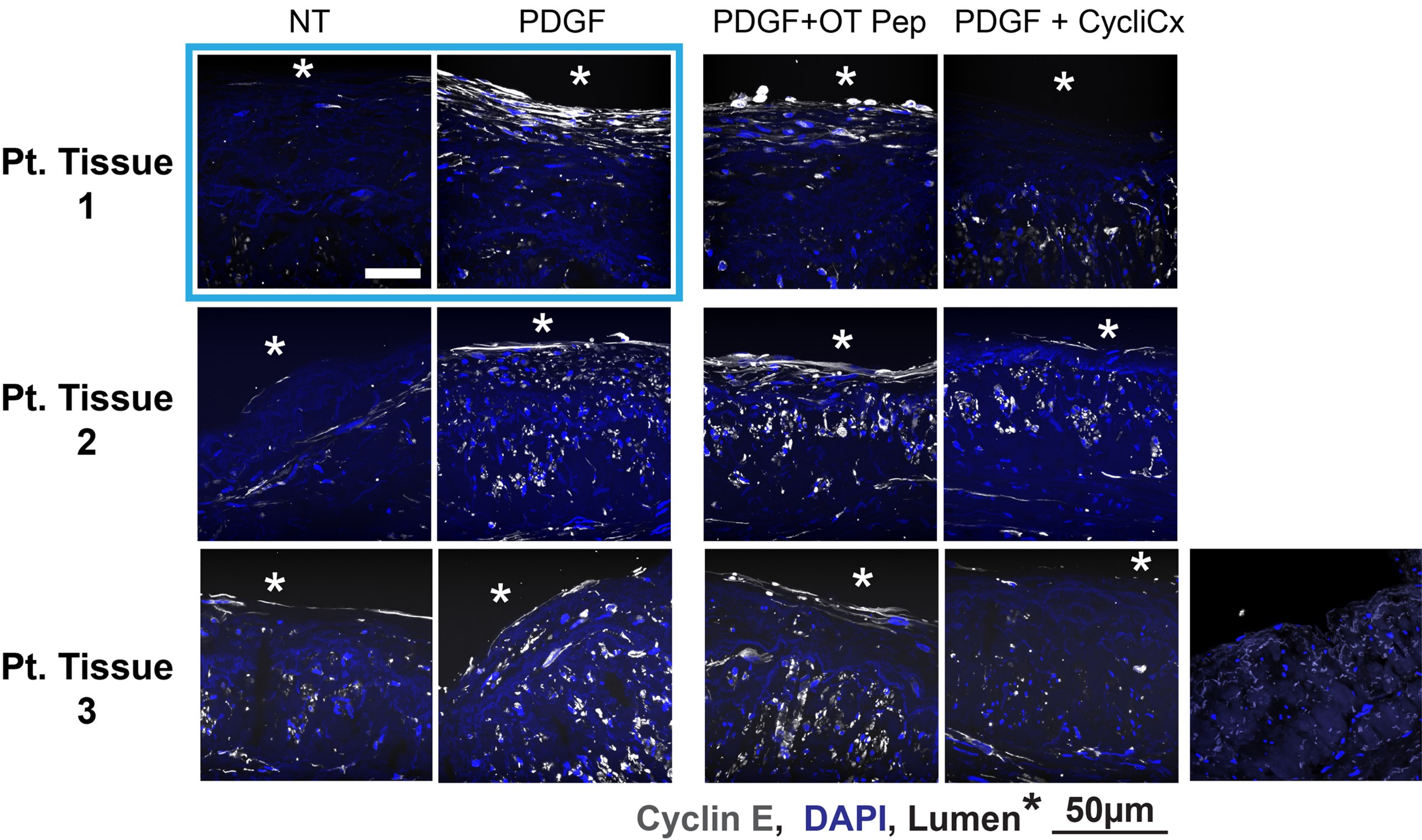

### Supplementary Figure 6: Peptide Array

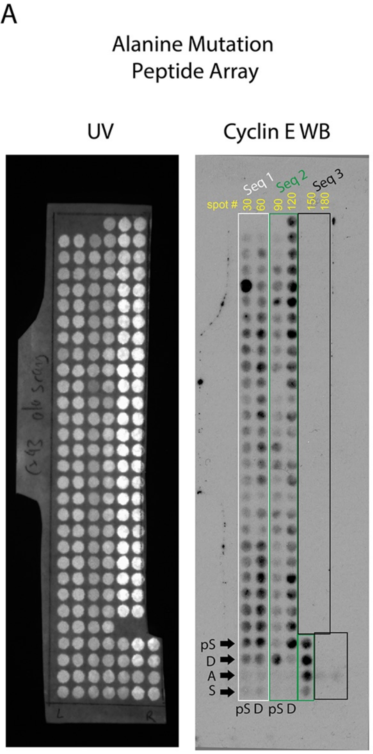


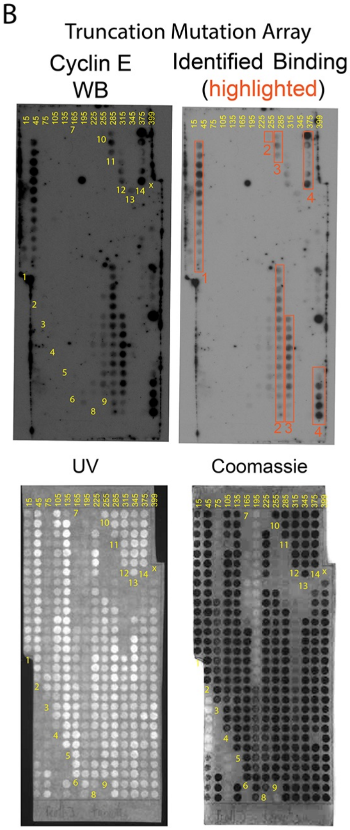


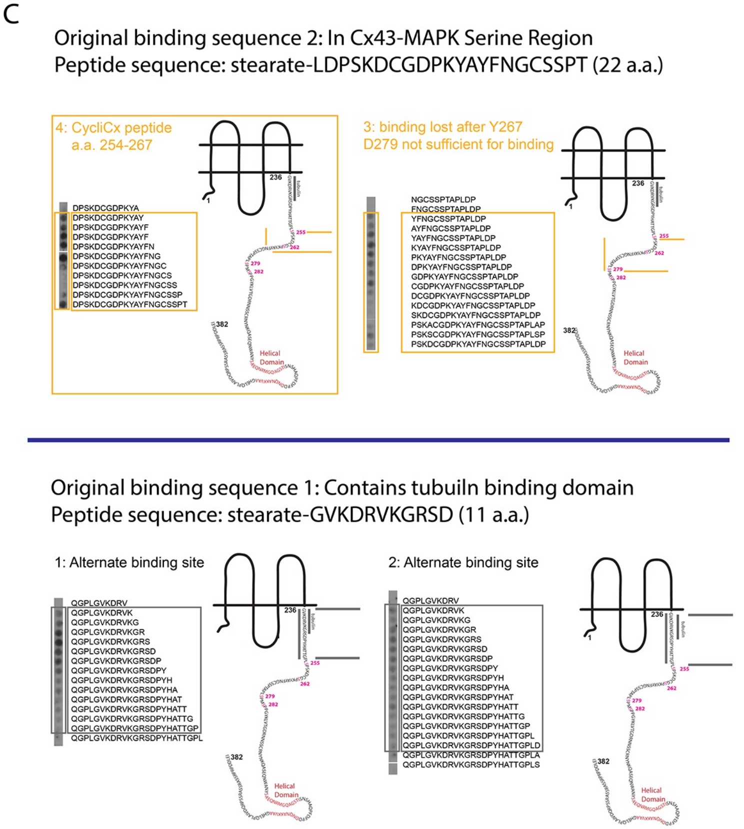


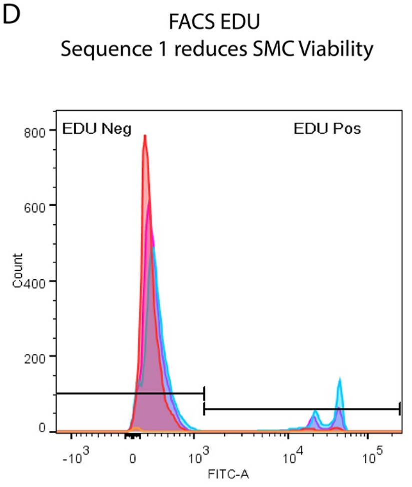


| | Cell # | EDU Neg (%) | EDU Pos (%) |
|---|---|---|---|
| NT | 10,000 | 89.8 | 10.2 |
| PDGF | 10,000 | 83.2 | 16.8 |
| PDGF + Pep1 | 106 | 98.1 | 1.89 |
| PDGF + Pep2 | 10,000 | 96.1 | 3.93 |

E

Final Peptide sequences:

| Peptide | sequence | Cx43 a.a. | a.a. | mw | charge |
|---|---|---|---|---|---|
| CycliCx (m) | stearate-LDPSKDCGDPKYAY | 254-267 | 14 | 1571 | -1 |
| CycliCx (h) | stearate-LDPAKDCGDQKYAY | 254-267 | 14 | 1586 | -1 |
| OT peptide | stearate-AYFNGCSSPTAPLDP | 265-280 | 15 | 1539 | -1 |

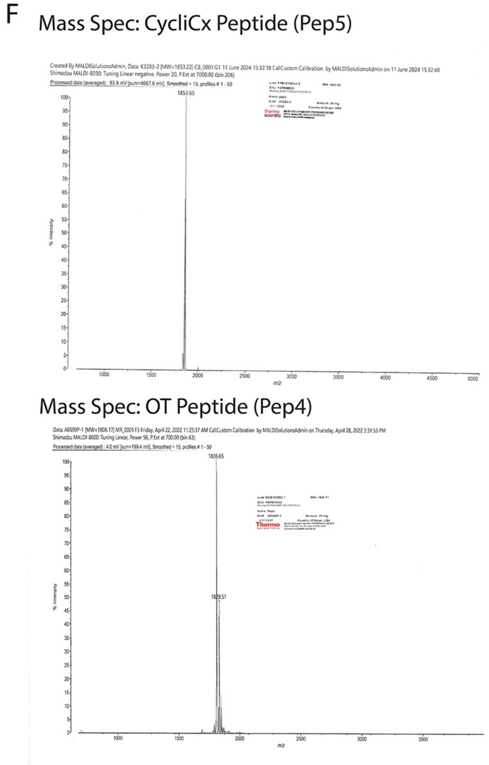

Supplemental Figure 07: Cx43 peptides do not alter PDGF-β-induced pERK in SMC

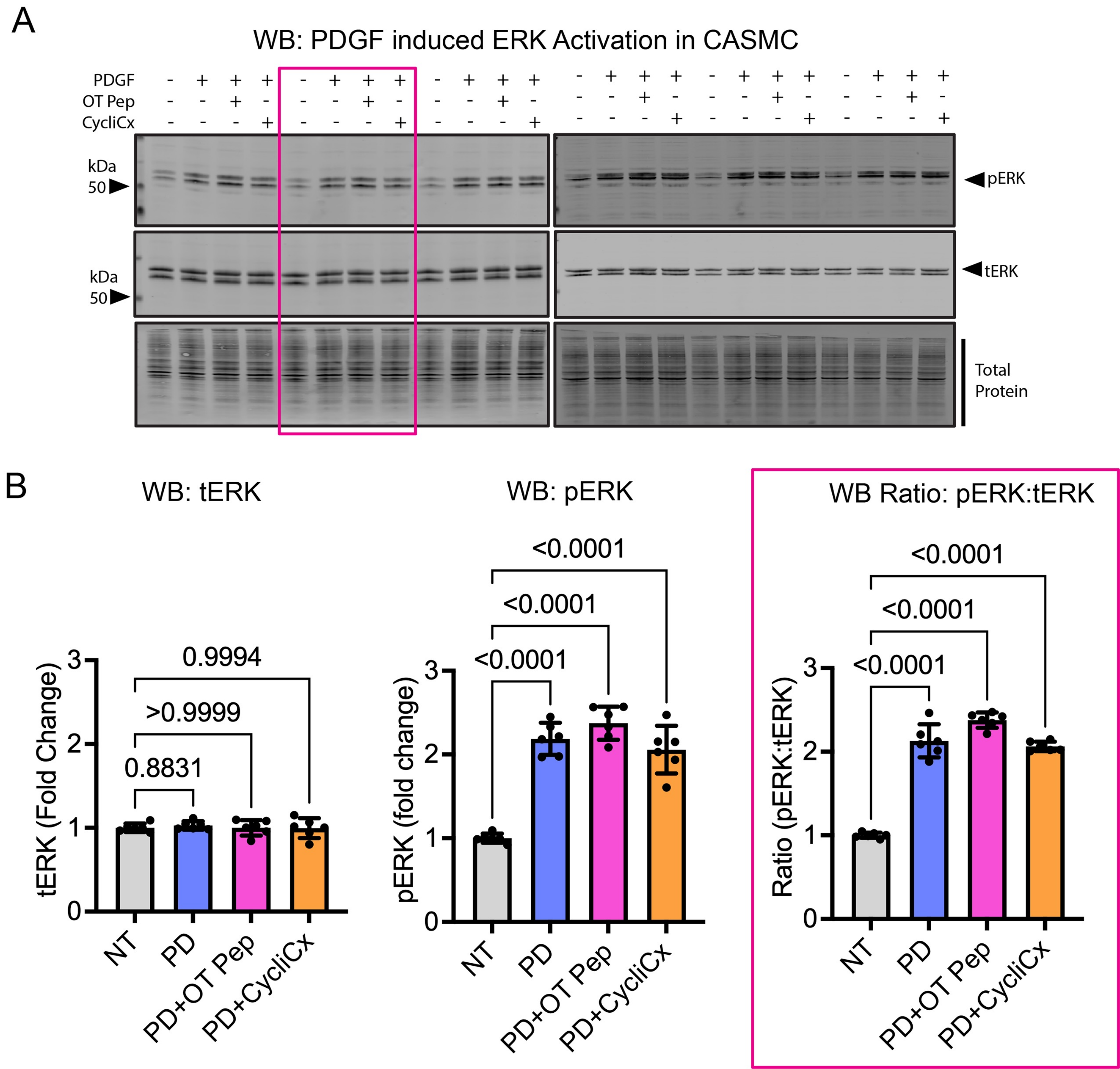

Supplemental Figure 08: CycliCx alters Cx43 phosphorylation

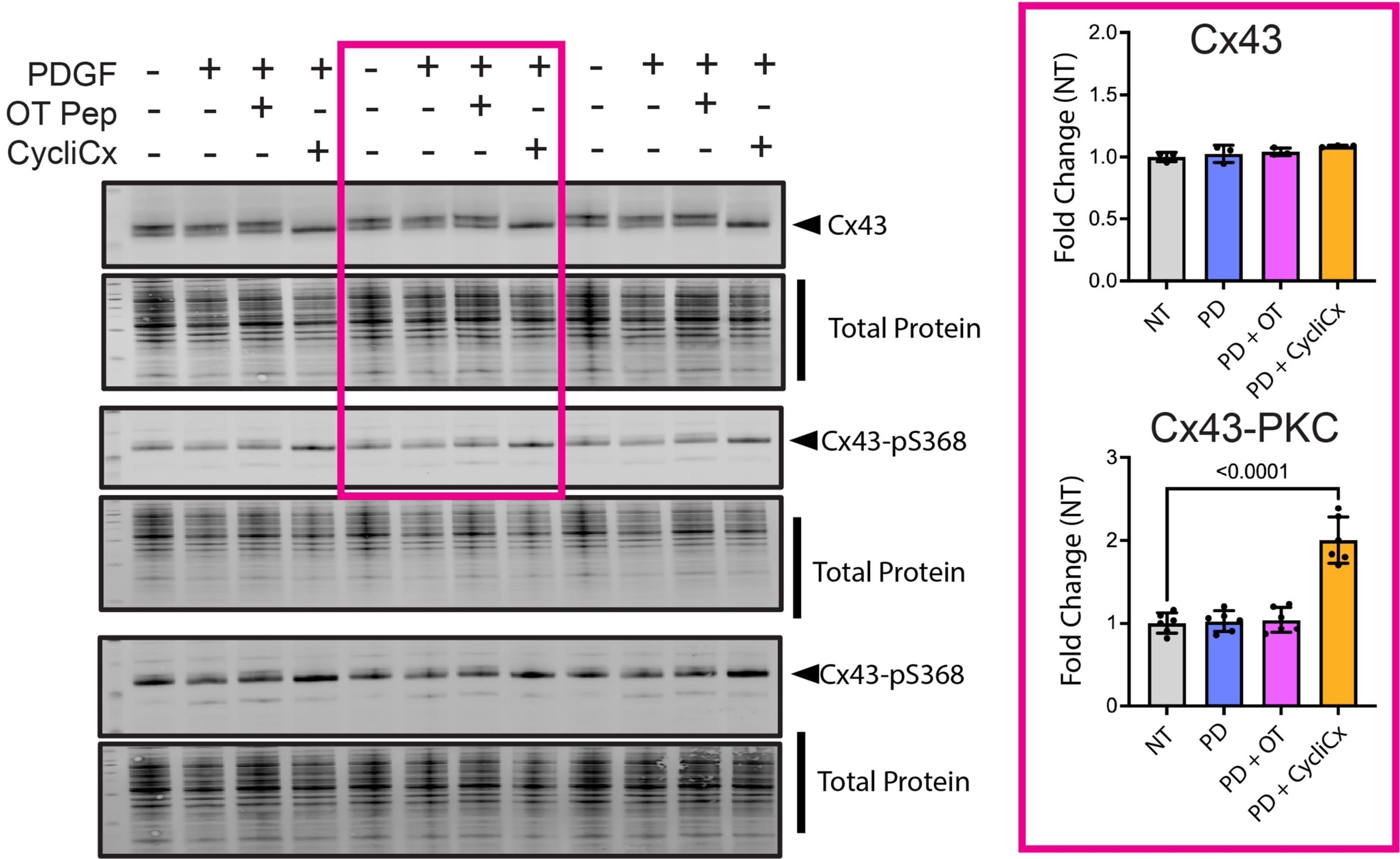

### Supplemental Figure 09: Timecourse, CycliCx alters Cx43 phosphorylation

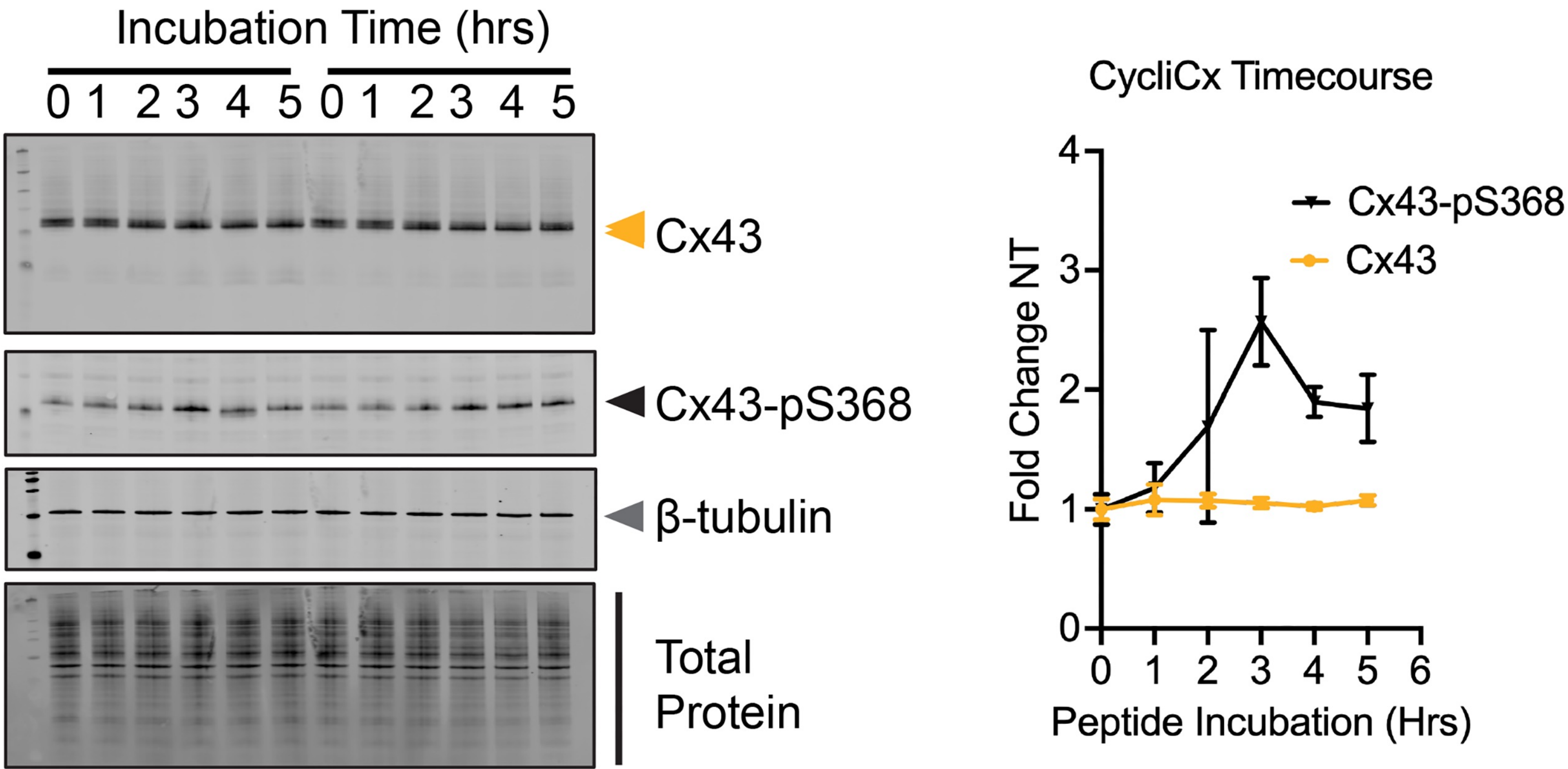

Supplemental Figure 10: IF Cx43 Peptide treatment CASMC

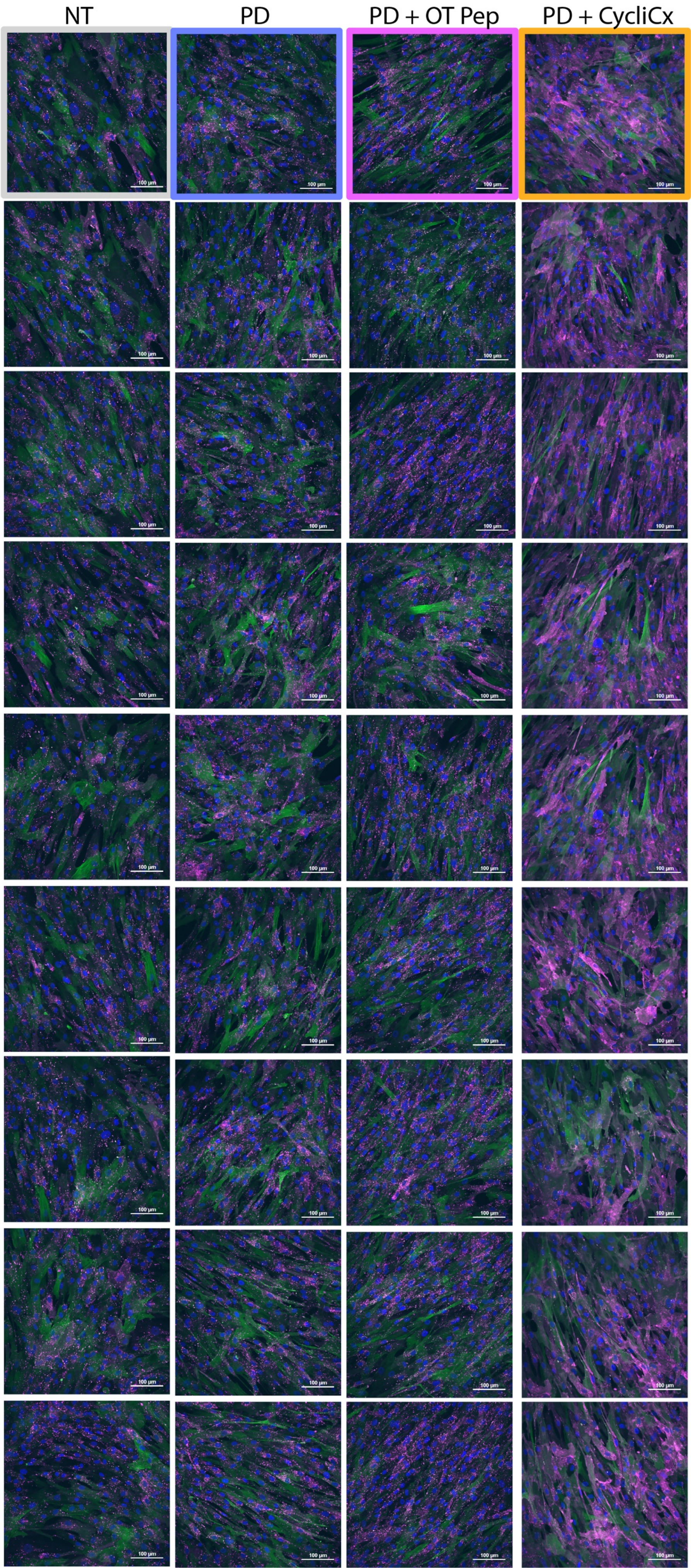

### Supplemental Figure 11: Cx43 Membrane Biotinylation

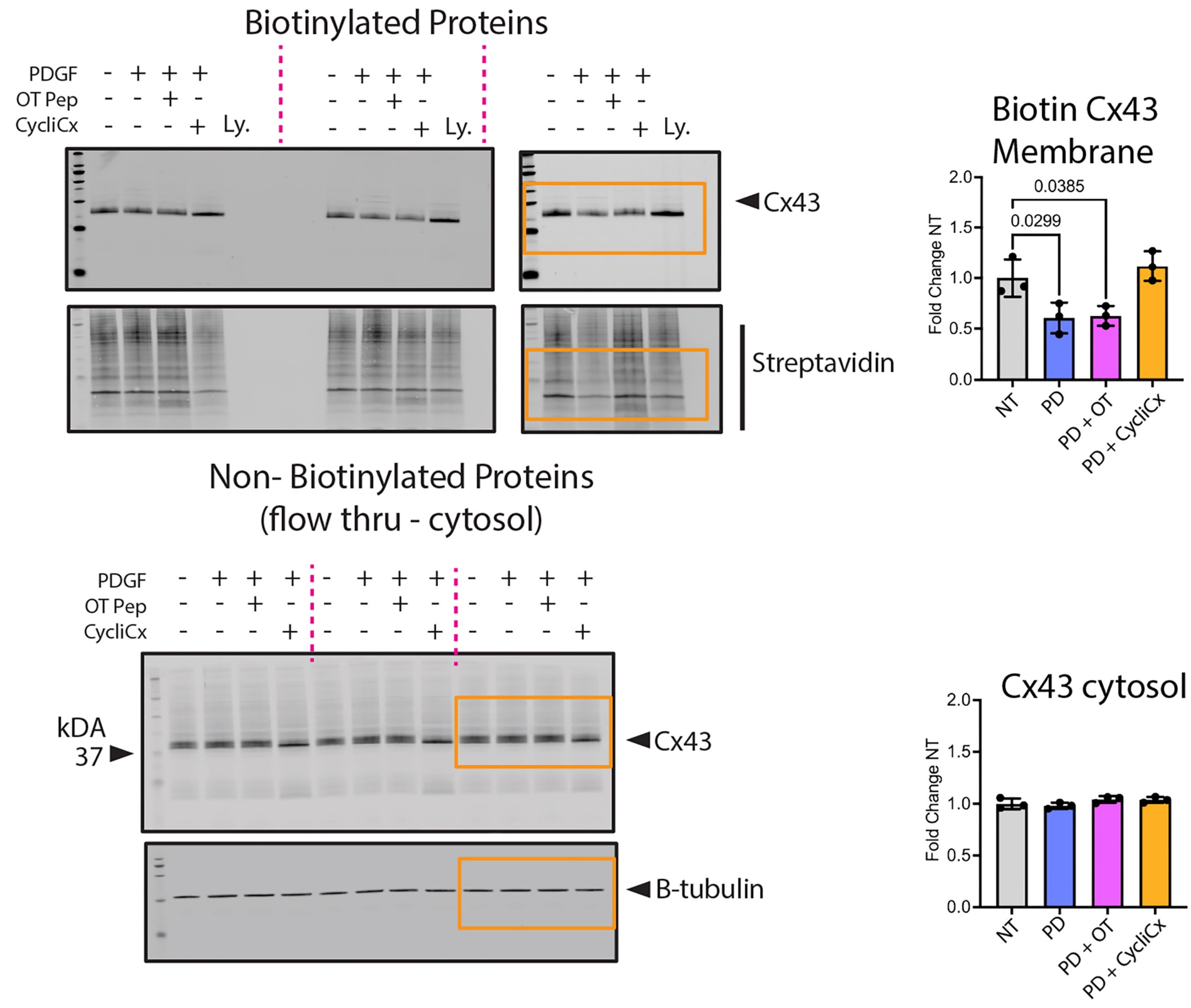

Supplemental Figure 12: Scrape load dye transfer (SLDT) and NIE image processing

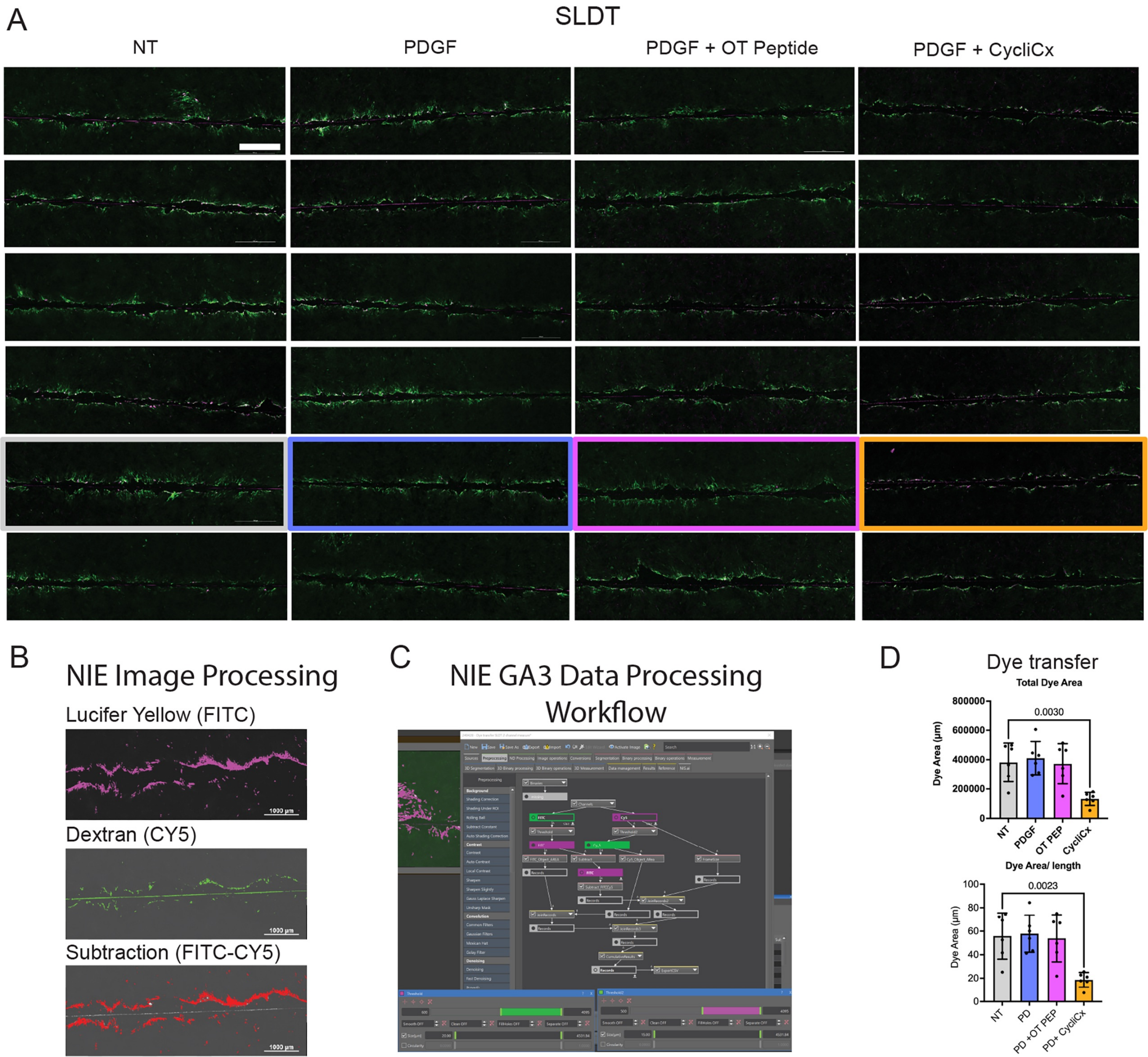

Supplemental Figure 13: Cx43 Co-immunoprecipitation in CASMC

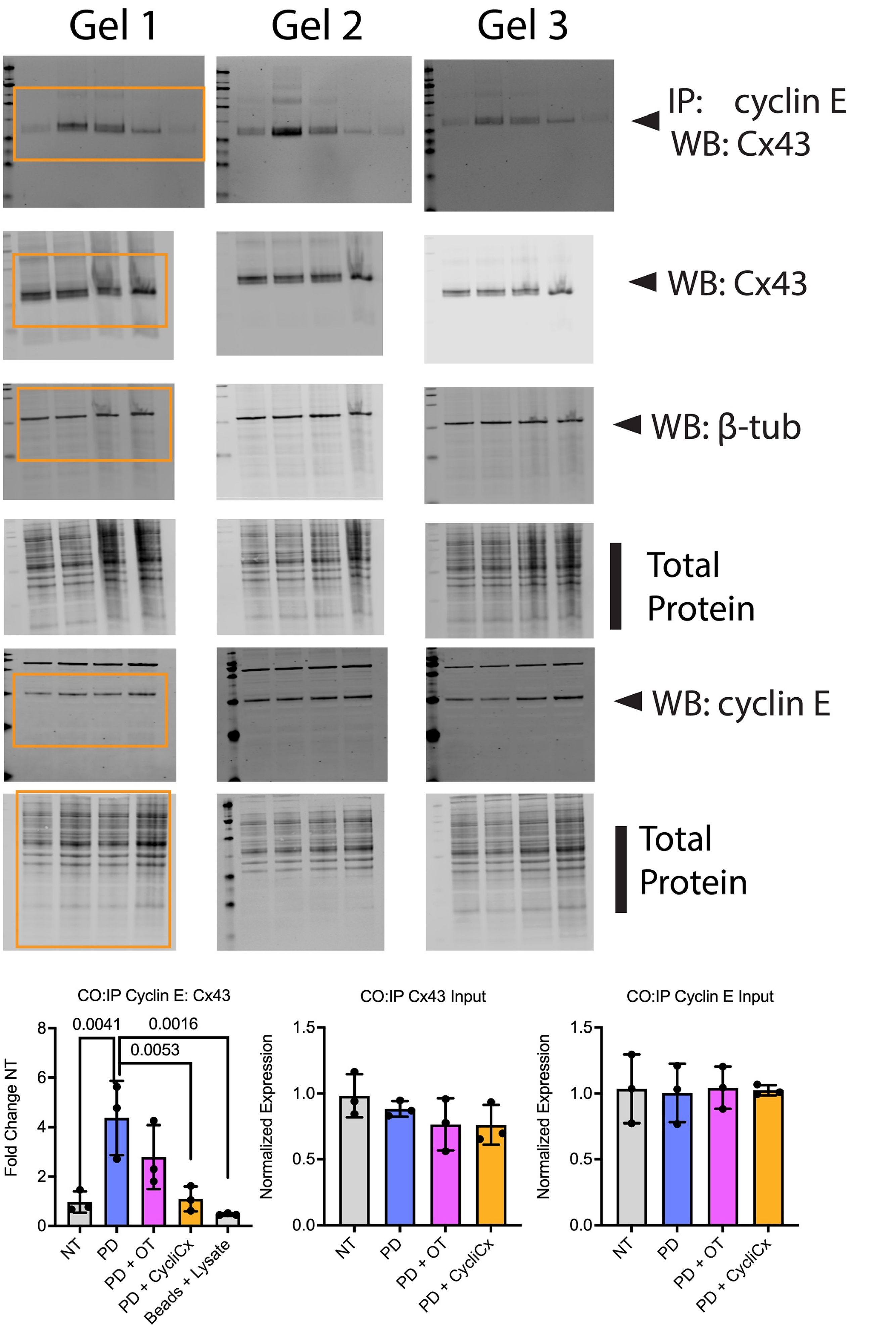

### Supplemental Figure 14: EDU Histogram Data

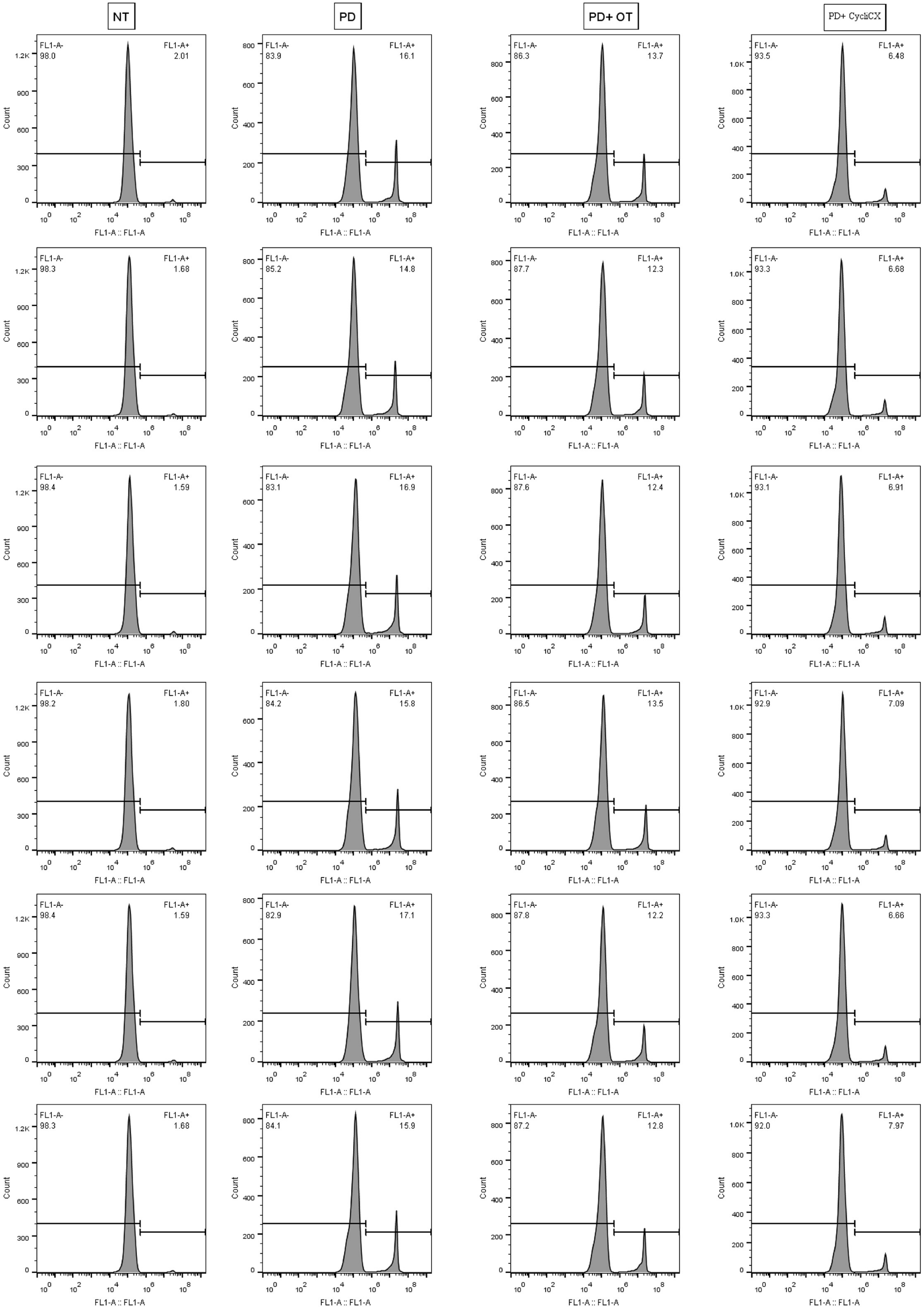

### Supplemental Figure 15: Bulk RNA sequencing quality control, correlation, and clustering

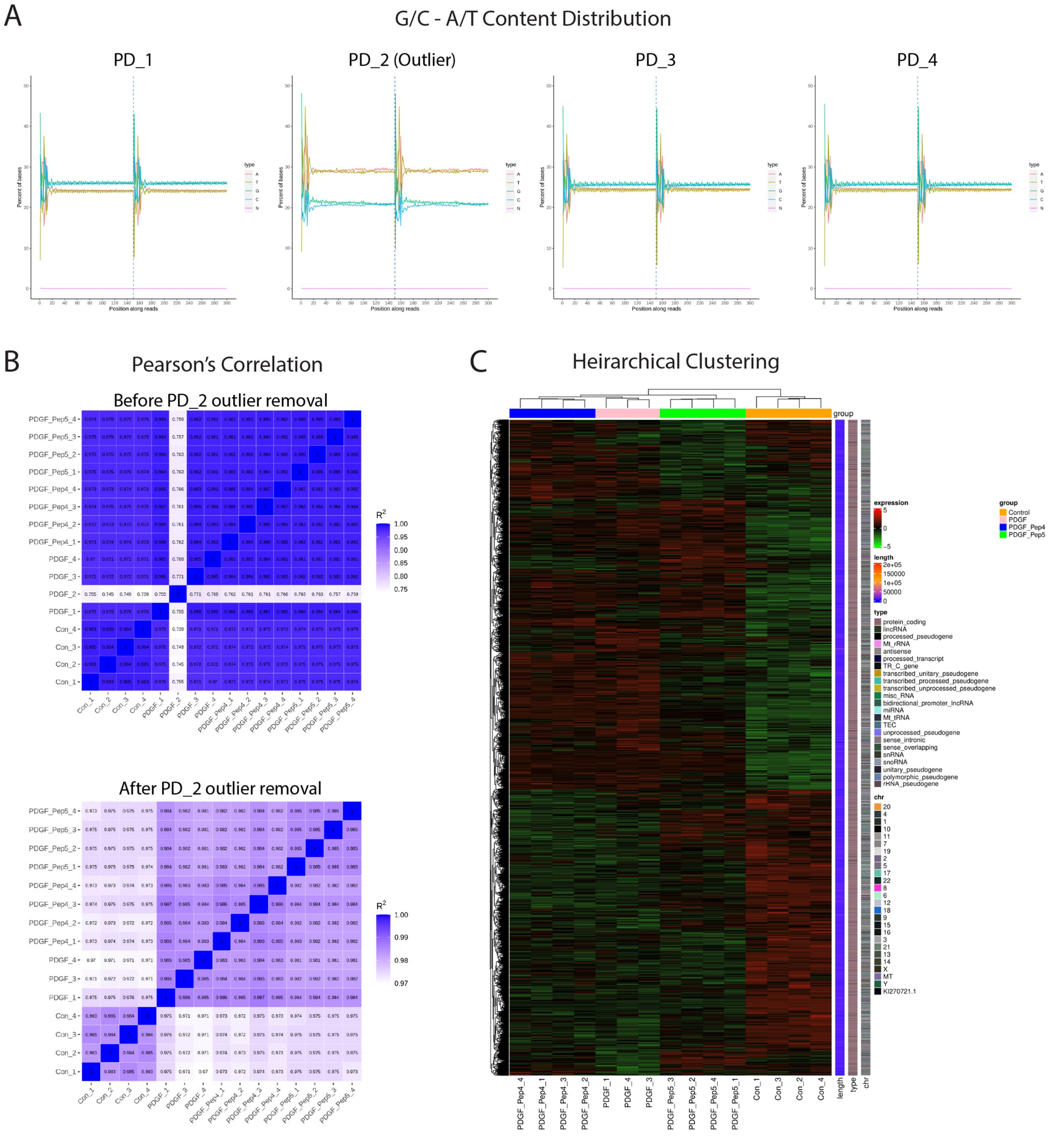

### Supplemental Figure 16: CycliCx peptide blunts mitotic gene pathways

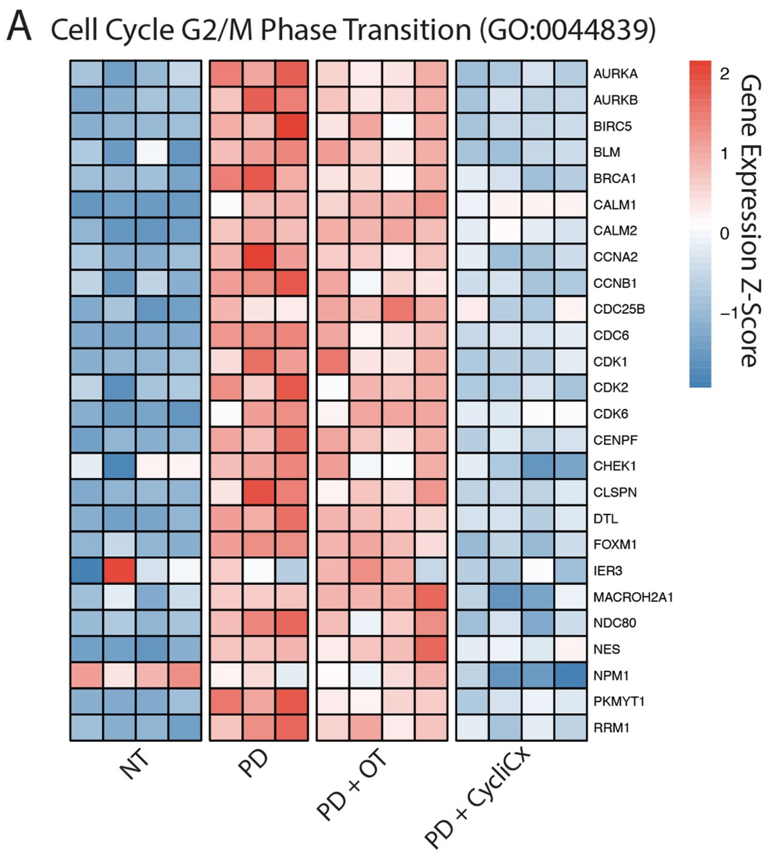


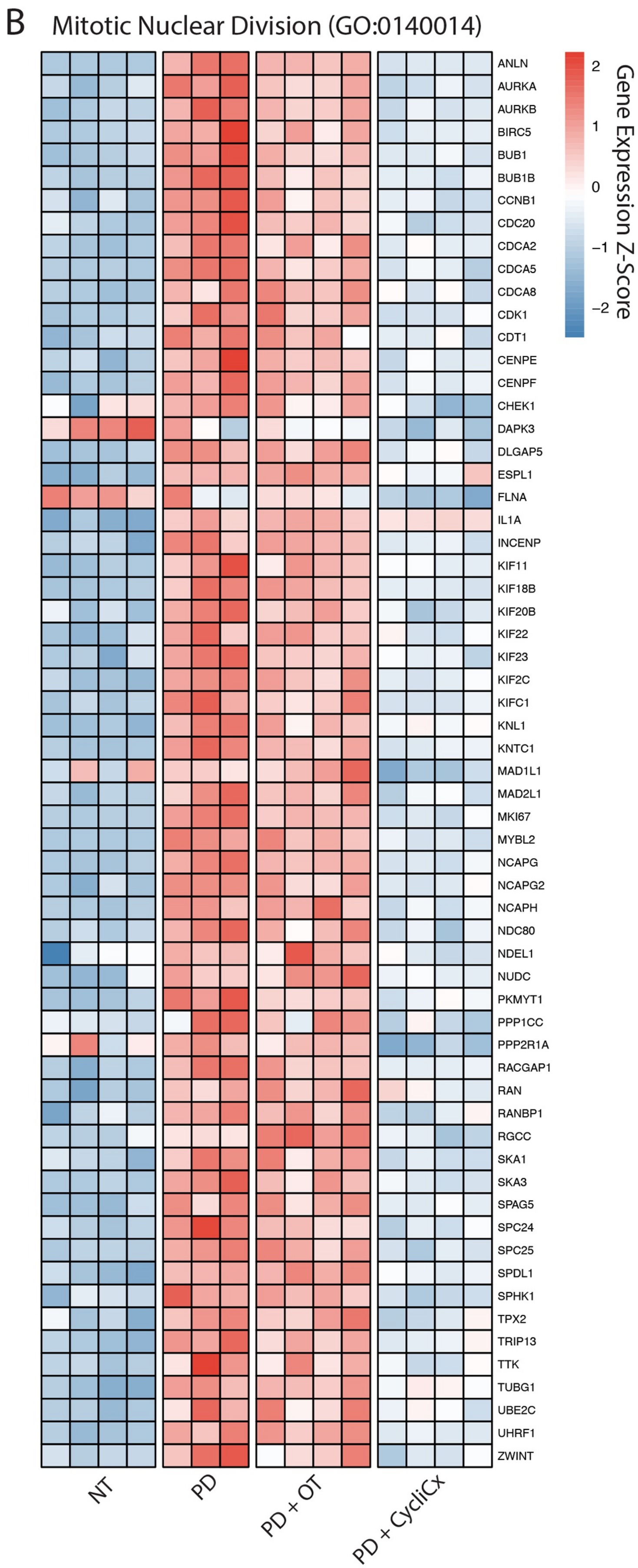

Supplemental Figure 17: CycliCx peptide alters the expression of critical cell cycle regulatory genes

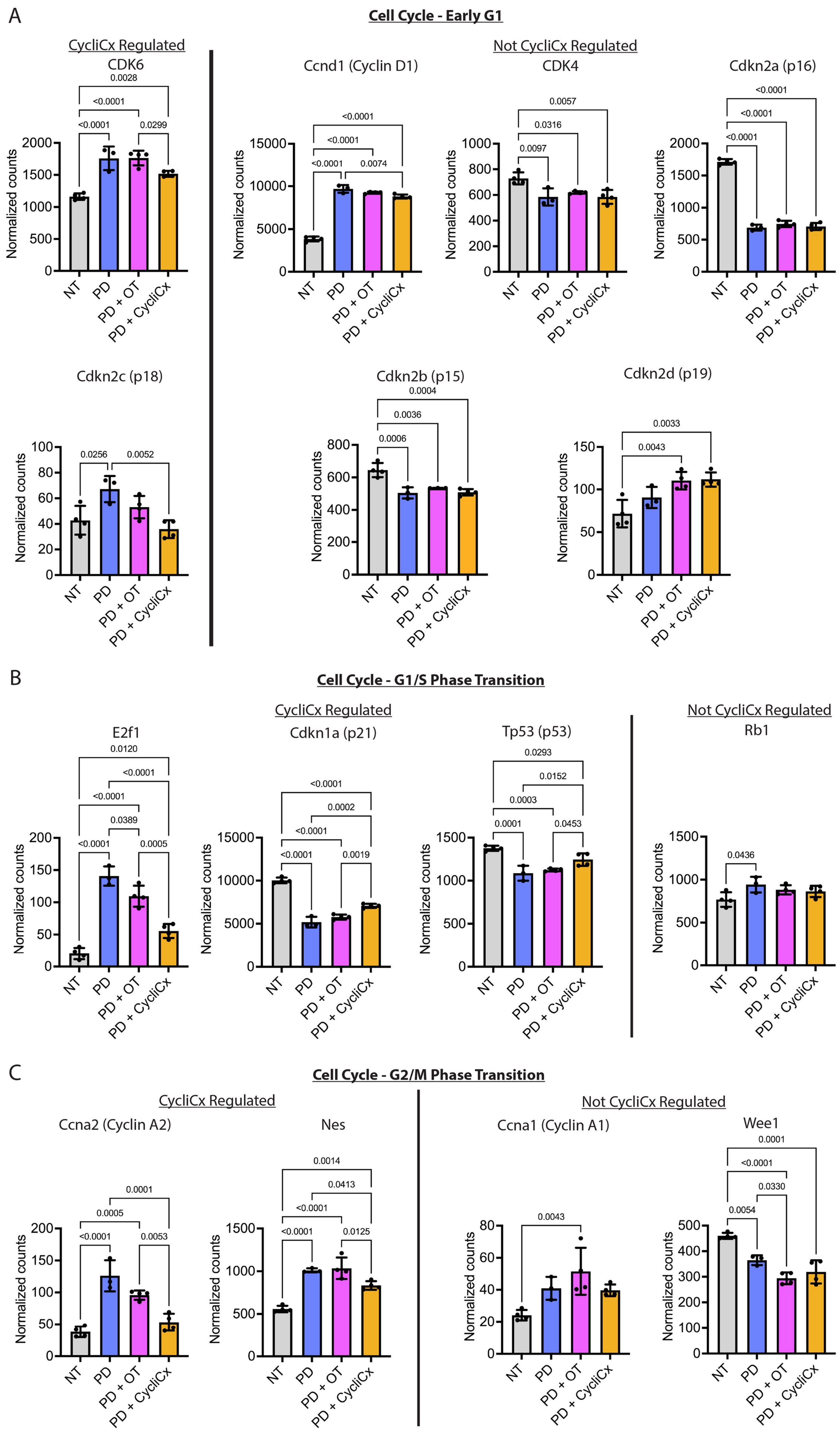

Supplemental Figure 18: Gene expression of gap junction proteins and PDGF regulators and transcription targets

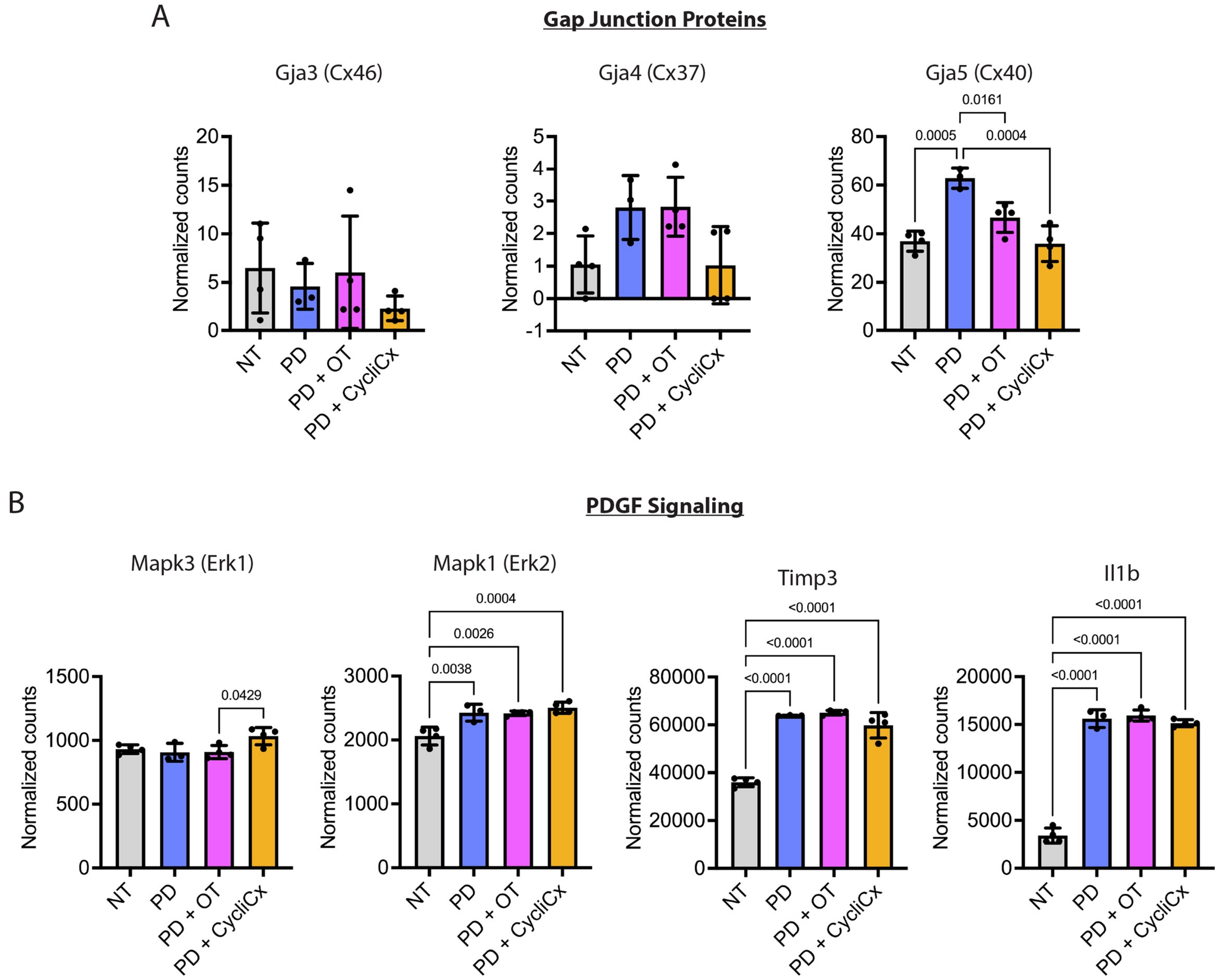

Supplemental Figure 19: CycliCx peptide blunts the expression of smooth muscle contractile genes

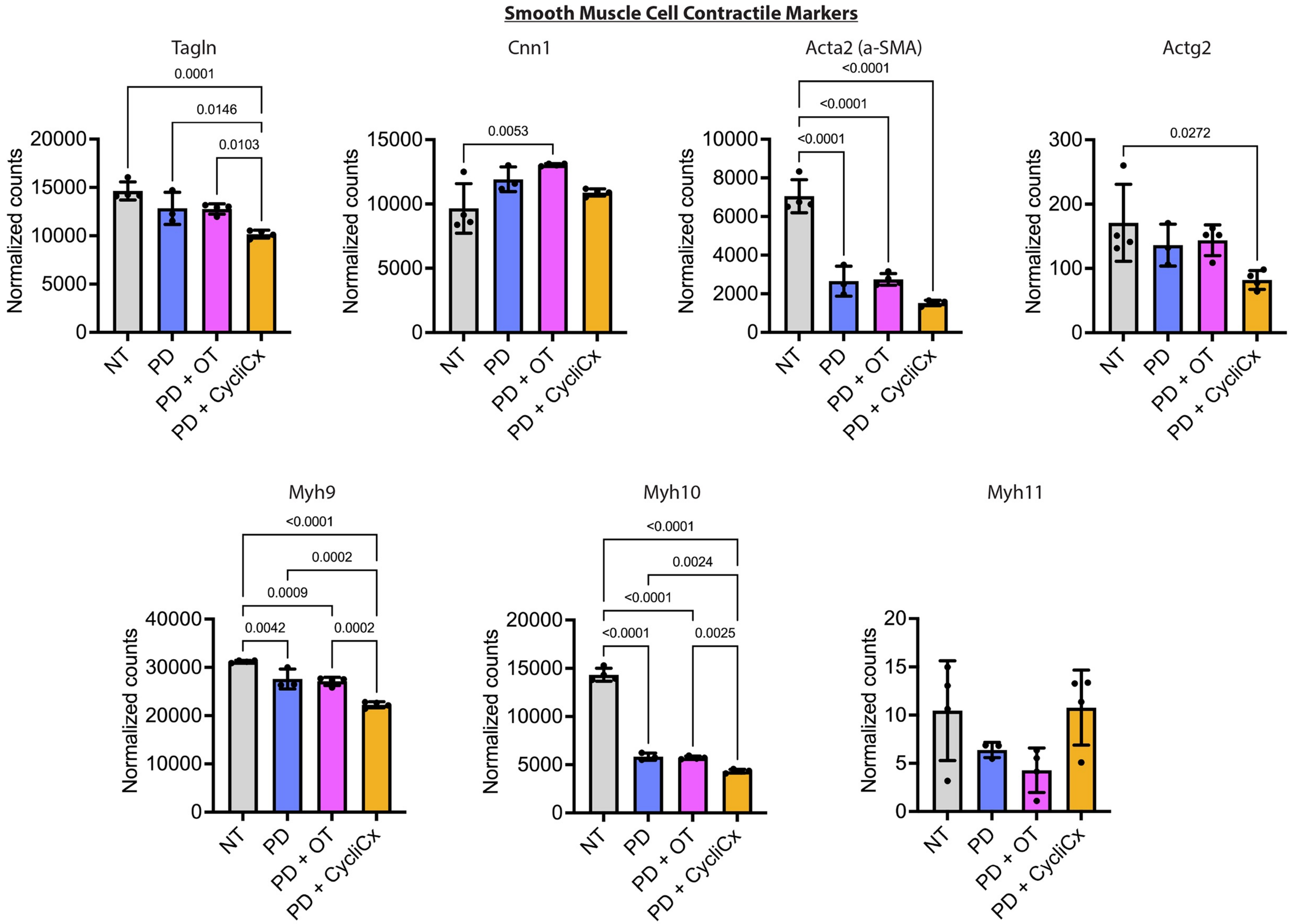

### Supplemental Figure 20: Mouse ligation neointima measurements

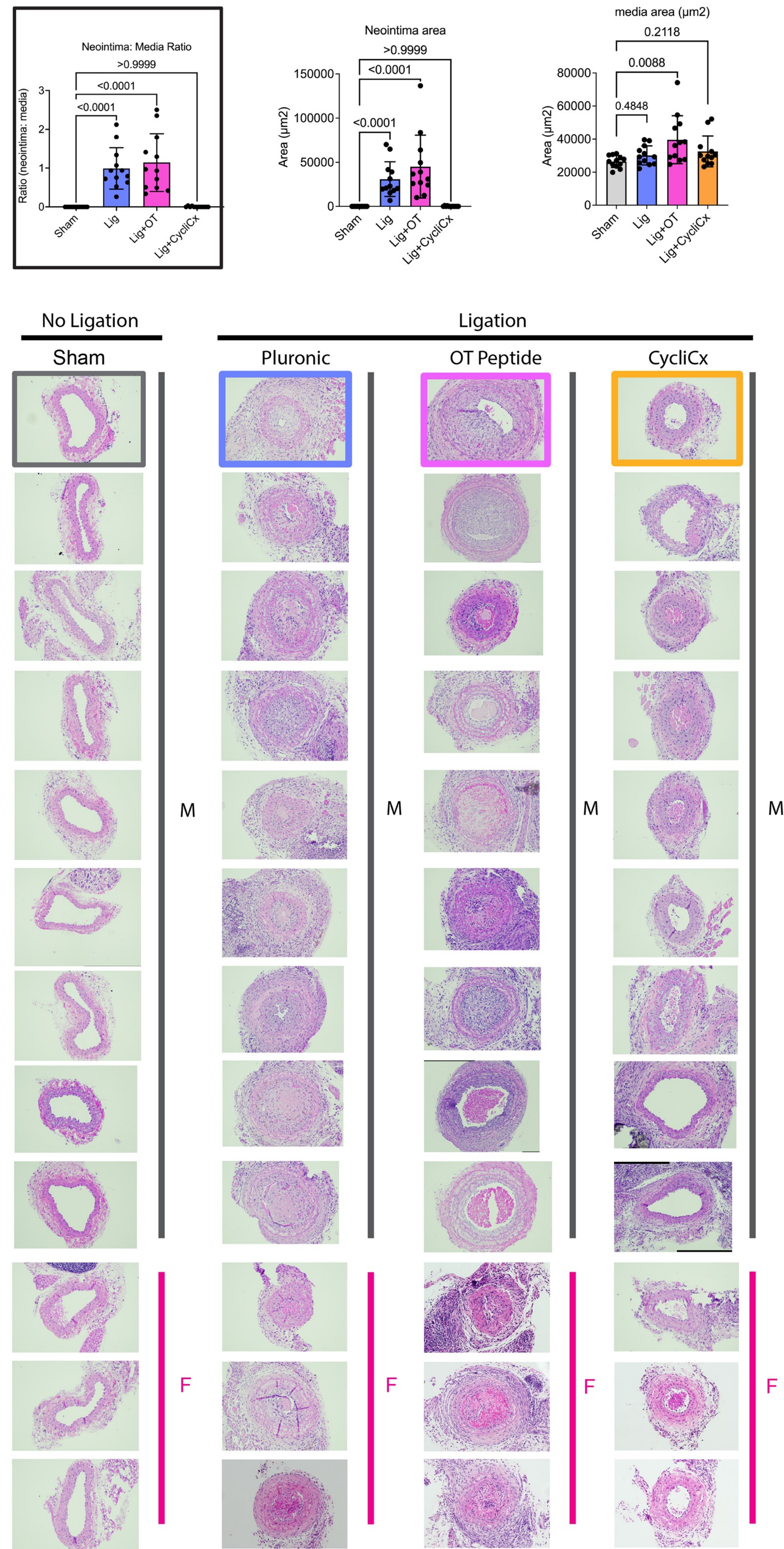
