## Supplementary material for "A connexin 43 targeting peptide prevents blood vessel neointima formation": Reagent list

**Supplemental Table 1: Key Resource Table**

| REAGENT or RESOURCE | SOURCE | IDENTIFIER |
|---|---|---|
| **Antibodies (IF/ WB/ CO-IP)** | | |
| Cx43 (WB/ CO-IP) | Millipore Sigma | C6219<br>RRID:AB_476857 |
| Cx43 (IF, Rabbit) | Millipore Sigma | HPA035097<br>UNSPSC:12352203 |
| Cx43 (WB/ CO-IP) | Millipore Sigma | C8093<br>RRID:AB_259085 |
| Phospho-Cx43 (pSer368) (WB) | Millipore Sigma | SAB4504371 |
| Phospho-Cx43 (pSer 279/282) 0.19 g/mL (IF) | custom antibody Source Dr. Paul Lampe, | Reference<br>PMID: 22652908 |
| Cyclin E1, CCNE1-HE12 (IF, Mouse) | Cell Signaling Technologies | 4129 |
| Cyclin E (CCNE1) (IF, WB, Rabbit) | Millipore Sigma | SAB1400044 |
| p44/42 MAPK (Erk1/2) (137F5, Rabbit) | Cell Signaling Technologies | 4695 |
| Phospho-p44/42 MAPK (Erk1/2) (Thr202/Tyr204) (D13.14.4E) XP®, Rabbit | Cell Signaling Technologies | 4370 |
| β-Tubulin (WB, Mouse) | DSHB | E7<br>RRID:AB_2315513 |
| Transgelin (IF, Goat) | Abcam | ab10135 |
| IRDye® 800CW, Donkey Anti-Rabbit | LI-COR Biosciences | 926-32213<br>RRID:AB_621848 |
| IRDye® 800CW, Donkey Anti-Mouse | LI-COR Biosciences | 926-32212,<br>RRID:AB_621847 |

| | | |
|---|---|---|
| IRDy(R) 680RD Donkey anti-Rabbit | LI-COR Biosciences | 926-68073 RRID:AB_10954442 |
| IRDye 680RD Donkey anti-Mouse | LI-COR Biosciences | 926-68072 RRID:AB_10953628 |
| IRDye®800CW Streptavidin | LI-COR Biosciences | 926-32230 |
| **Chemicals and Reagents** | | |
| Gibco™ M231 media | ThermoFisher Scientific | M-231-500 |
| VWR Life Science Seradigm Premium Grade Fetal Bovine Serum | VWR Life Science | 97068-085 |
| Smooth muscle growth supplement (SMGS) | ThermoFisher Scientific | S00725 |
| Trypsin-EDTA (0.5%), no phenol red | ThermoFisher Scientific | 15400-054 |
| Gibco™ M231 media | ThermoFisher Scientific | M-231-500 |
| DPBS (1X), no calcium, no magnesium | ATCC | 30-2200 |
| Recombinant PDGF-bb | Millipore | 01-305 |
| Ethylene glycol-bis(2-aminoethylether)-N,N,N′,N′-tetraacetic acid | Sigma-Aldrich | E4378 |
| Protease Inhibitor Cocktail | Millipore Sigma | P8340 |
| phosphatase Inhibitor Cocktail 2 | Millipore Sigma | P5726 |
| phosphatase Inhibitor Cocktail 3 | Millipore Sigma | P0044 |
| RNeasy kit | Qiagen | 74004 |
| Odyssey Blocking Buffer (PBS) | LI-COR Biosciences | 927-40000 |

| | | |
|---|---|---|
| Dynabeads™ M-280 Streptavidin | ThermoFisher Scientific | 11205D |
| EZ-Link™ Sulfo-NHS-LC-Biotin | ThermoFisher Scientific | 21335 |
| 5-ethynyl-2'deoxyuridine (EDU) Flow cytometry kitt-488 ( | ThermoFisher Scientific | C10425 |
| Ethiqa XR 1.3 mg/mL | Covetrus | 072117 |
| Isoflurane | Covetrus | 029405 |
| Pluronic F127 | Millipore Sigma | P2443 |
| CycliCx Peptide: stearate-LDPSKDCGDPKYAY | ThermoFisher custom peptide synthesis services | Custom |
| OT Peptide: stearate-AYFNGCSSPTAPLDP | ThermoFisher custom peptide synthesis services | Custom |
| **Commercial Assays** | | |
| Pierce™ BCA Protein Assay Kit | ThermoFisher Scientific | 23225 |
| REVERT™ Total Protein Stain for immunoblot Normalization | LI-COR Biosciences | 926-11010 |
| **Experimental Models: Cell lines and Mice** | | |
| CASMC: primary human SMC | ATCC | PCS-100-02 |
| CASMC: primary human SMC | Lonza | CC-2583 |
| CASMC: primary human SMC | Thermo Fisher Scientific | C0175C |
| C57BL/6J mice | Jackson Laboratory | 000664 |

**Key:**
IF – Immunofluorescence

WB – Western blot
Co-IP co-immunoprecipitation
