## Supplementary material for "A connexin 43 targeting peptide prevents blood vessel neointima formation": Peptide array sequence list

Supplemental List Cx43CT Truncation Mutation array
Full sequence
QGPLGVKDRVKGRSDPYHATTGPLSPSKDCGSPKYAYFNGCSSPTAPLSPMSPPGYKLVTGDRNNSSCRN

Arrays

(1)

1. Q G P L G V K D R V K G R S D P Y H A T T G P L pS
2. Q G P L G V K D R V K G R S D P Y H A T T G P L S
3. Q G P L G V K D R V K G R S D P Y H A T T G P L A
4. G P L G V K D R V K G R S D P Y H A T T G P L pS
5. P L G V K D R V K G R S D P Y H A T T G P L pS
6. L G V K D R V K G R S D P Y H A T T G P L pS
7. G V K D R V K G R S D P Y H A T T G P L pS
8. V K D R V K G R S D P Y H A T T G P L pS
9. K D R V K G R S D P Y H A T T G P L pS
10. D R V K G R S D P Y H A T T G P L pS
11. R V K G R S D P Y H A T T G P L pS
12. V K G R S D P Y H A T T G P L pS
13. K G R S D P Y H A T T G P L pS
14. G R S D P Y H A T T G P L pS
15. R S D P Y H A T T G P L pS
16. S D P Y H A T T G P L pS
17. D P Y H A T T G P L pS
18. P Y H A T T G P L pS
19. Y H A T T G P L pS
20. H A T T G P L pS
21. A T T G P L pS
22. T T G P L pS
23. T G P L pS
24. P L pS
25. L pS
26. pS
27. .space

(2)

1. Q G P L G V K D R V K G R S D P Y H A T T G P L pS
2. Q G P L G V K D R V K G R S D P Y H A T T G P L S
3. Q G P L G V K D R V K G R S D P Y H A T T G P L A
4. Q G P L G V K D R V K G R S D P Y H A T T G P L
5. Q G P L G V K D R V K G R S D P Y H A T T G P
6. Q G P L G V K D R V K G R S D P Y H A T T G
7. Q G P L G V K D R V K G R S D P Y H A T T
8. Q G P L G V K D R V K G R S D P Y H A T
9. Q G P L G V K D R V K G R S D P Y H A
10. Q G P L G V K D R V K G R S D P Y H
11. Q G P L G V K D R V K G R S D P Y
12. Q G P L G V K D R V K G R S D P
13. Q G P L G V K D R V K G R S D
14. Q G P L G V K D R V K G R S
15. Q G P L G V K D R V K G R
16. Q G P L G V K D R V K G
17. Q G P L G V K D R V K
18. Q G P L G V K D R V
19. Q G P L G V K D R
20. Q G P L G V K D
21. Q G P L G V K
22. Q G P L G V
23. Q G P L G
24. Q G P L
25. Q G P
26. Q G
27. Q
28. .space

(3)pS

1. P S K D C G pS P K Y A Y F N G C S S P T A P L pS P
2. P S K D C G S P K Y A Y F N G C S S P T A P L S P
3. P S K D C G A P K Y A Y F N G C S S P T A P L A P
4. S K D C G pS P K Y A Y F N G C S S P T A P L pS P
5. K D C G pS P K Y A Y F N G C S S P T A P L pS P
6. D C G pS P K Y A Y F N G C S S P T A P L pS P
7. C G pS P K Y A Y F N G C S S P T A P L pS P

8. G pS P K Y A Y F N G C S S P T A P L pS P
9. pS P K Y A Y F N G C S S P T A P L pS P
10. P K Y A Y F N G C S S P T A P L pS P
11. K Y A Y F N G C S S P T A P L pS P
12. Y A Y F N G C S S P T A P L pS P
13. A Y F N G C S S P T A P L pS P
14. Y F N G C S S P T A P L pS P
15. F N G C S S P T A P L pS P
16. N G C S S P T A P L pS P
17. G C S S P T A P L pS P
18. C S S P T A P L pS P
19. S S P T A P L pS P
20. S P T A P L pS P
21. P T A P L pS P
22. A P L pS P
23. P L pS P
24. L pS P
25. pS P
26. P
27. .space

(4)pS

1. P S K D C G pS P K Y A Y F N G C S S P T A P L pS P
2. P S K D C G S P K Y A Y F N G C S S P T A P L S P
3. P S K D C G A P K Y A Y F N G C S S P T A P L A P
4. P S K D C G pS P K Y A Y F N G C S S P T A P L pS
5. P S K D C G pS P K Y A Y F N G C S S P T A P L
6. P S K D C G pS P K Y A Y F N G C S S P T A P
7. P S K D C G pS P K Y A Y F N G C S S P T A
8. P S K D C G pS P K Y A Y F N G C S S P T
9. P S K D C G pS P K Y A Y F N G C S S P
10. P S K D C G pS P K Y A Y F N G C S S
11. P S K D C G pS P K Y A Y F N G C S
12. P S K D C G pS P K Y A Y F N G C
13. P S K D C G pS P K Y A Y F N G
14. P S K D C G pS P K Y A Y F N
15. P S K D C G pS P K Y A Y F
16. P S K D C G pS P K Y A Y
17. P S K D C G pS P K Y A
18. P S K D C G pS P K Y
19. P S K D C G pS P K
20. P S K D C G pS P
21. P S K D C G pS
22. P S K D C G
23. P S K D C
24. P S K D
25. P S K
26. P S
27. P
28. .space

(5)

1. A P L pS P M pS P P G Y K L V T G D R N N S S C R N
2. A P L S P M S P P G Y K L V T G D R N N S S C R N
3. A P L A P M A P P G Y K L V T G D R N N S S C R N
4. P L pS P M pS P P G Y K L V T G D R N N S S C R N
5. P L pS P M pS P P G Y K L V T G D R N N S S C R N
6. pS P M pS P P G Y K L V T G D R N N S S C R N
7. P M pS P P G Y K L V T G D R N N S S C R N
8. M pS P P G Y K L V T G D R N N S S C R N
9. pS P P G Y K L V T G D R N N S S C R N
10. P P G Y K L V T G D R N N S S C R N
11. P G Y K L V T G D R N N S S C R N
12. G Y K L V T G D R N N S S C R N
13. Y K L V T G D R N N S S C R N
14. K L V T G D R N N S S C R N
15. L V T G D R N N S S C R N
16. V T G D R N N S S C R N
17. T G D R N N S S C R N
18. G D R N N S S C R N
19. R N N S S C R N
20. N N S S C R N
21. N S S C R N

22. S S C R N
23. S C R N
24. C R N
25. R N
26. N
27. .space

(6) pS

1. A P L pS P M pS P P G Y K L V T G D R N N S S C R N
2. A P L S P M S P P G Y K L V T G D R N N S S C R N
3. A P L A P M A P P G Y K L V T G D R N N S S C R N
4. A P L pS P M pS P P G Y K L V T G D R N N S S C R
5. A P L pS P M pS P P G Y K L V T G D R N N S S C
6. A P L pS P M pS P P G Y K L V T G D R N N S S
7. A P L pS P M pS P P G Y K L V T G D R N N S
8. A P L pS P M pS P P G Y K L V T G D R N N
9. A P L pS P M pS P P G Y K L V T G D R N
10. A P L pS P M pS P P G Y K L V T G D R
11. A P L pS P M pS P P G Y K L V T G D
12. A P L pS P M pS P P G Y K L V T G
13. A P L pS P M pS P P G Y K L V T
14. A P L pS P M pS P P G Y K L V
15. A P L pS P M pS P P G Y K L
16. A P L pS P M pS P P G Y K
17. A P L pS P M pS P P G Y
18. A P L pS P M pS P P G
19. A P L pS P M pS P P
20. A P L pS P M pS P
21. A P L pS P M pS
22. A P L pS P M
23. A P L pS P
24. A P L pS
25. A P L
26. A P
27. A
28. .space

(7) pS

1. pS P S K D C G pS P K Y A Y F N G C S S P T A P L pS P M pS
2. S P S K D C G S P K Y A Y F N G C S S P T A P L S P M S
3. A P S K D C G A P K Y A Y F N G C S S P T A P L A P M A
4. P S K D C G pS P K Y A Y F N G C S S P T A P L pS P M pS
5. S K D C G pS P K Y A Y F N G C S S P T A P L pS P M pS
6. K D C G pS P K Y A Y F N G C S S P T A P L pS P M pS
7. D C G pS P K Y A Y F N G C S S P T A P L pS P M pS
8. C G pS P K Y A Y F N G C S S P T A P L pS P M pS
9. G pS P K Y A Y F N G C S S P T A P L pS P M pS
10. pS P K Y A Y F N G C S S P T A P L pS P M pS
11. P K Y A Y F N G C S S P T A P L pS P M pS
12. K Y A Y F N G C S S P T A P L pS P M pS
13. Y A Y F N G C S S P T A P L pS P M pS
14. A Y F N G C S S P T A P L pS P M pS
15. Y F N G C S S P T A P L pS P M pS
16. F N G C S S P T A P L pS P M pS
17. N G C S S P T A P L pS P M pS
18. G C S S P T A P L pS P M pS
19. C S S P T A P L pS P M pS
20. S S P T A P L pS P M pS
21. S P T A P L pS P M pS
22. P T A P L pS P M pS
23. T A P L pS P M pS
24. A P L pS P M pS
25. P L pS P M pS
26. L pS P M pS
27. pS P M pS
28. P M pS
29. M pS
30. pS
31. .space

(8) pS

1. pS P S K D C G pS P K Y A Y F N G C S S P T A P L pS P M pS

2. S P S K D C G S P K Y A Y F N G C S S P T A P L S P M S
3. A P S K D C G A P K Y A Y F N G C S S P T A P L A P M A
4. pS P S K D C G pS P K Y A Y F N G C S S P T A P L pS P M
5. pS P S K D C G pS P K Y A Y F N G C S S P T A P L pS P
6. pS P S K D C G pS P K Y A Y F N G C S S P T A P L pS
7. pS P S K D C G pS P K Y A Y F N G C S S P T A P L
8. pS P S K D C G pS P K Y A Y F N G C S S P T A P
9. pS P S K D C G pS P K Y A Y F N G C S S P T A
10. pS P S K D C G pS P K Y A Y F N G C S S P T
11. pS P S K D C G pS P K Y A Y F N G C S S P
12. pS P S K D C G pS P K Y A Y F N G C S S
13. pS P S K D C G pS P K Y A Y F N G C S
14. pS P S K D C G pS P K Y A Y F N G C
15. pS P S K D C G pS P K Y A Y F N G
16. pS P S K D C G pS P K Y A Y F N
17. pS P S K D C G pS P K Y A Y F
18. pS P S K D C G pS P K Y A Y
19. pS P S K D C G pS P K Y A
20. pS P S K D C G pS P K Y
21. pS P S K D C G pS P K
22. pS P S K D C G pS P
23. pS P S K D C G pS
24. pS P S K D C G
25. pS P S K D C
26. pS P S K D
27. pS P S K
28. pS P S
29. pS P
30. pS
31. sp

(9) D

1. Q G P L G V K D R V K G R S D P Y H A T T G P L D
2. Q G P L G V K D R V K G R S D P Y H A T T G P L S
3. Q G P L G V K D R V K G R S D P Y H A T T G P L A
4. G P L G V K D R V K G R S D P Y H A T T G P L D
5. P L G V K D R V K G R S D P Y H A T T G P L D
6. L G V K D R V K G R S D P Y H A T T G P L D
7. G V K D R V K G R S D P Y H A T T G P L D
8. V K D R V K G R S D P Y H A T T G P L D
9. K D R V K G R S D P Y H A T T G P L D
10. D R V K G R S D P Y H A T T G P L D
11. R V K G R S D P Y H A T T G P L D
12. V K G R S D P Y H A T T G P L D
13. K G R S D P Y H A T T G P L D
14. G R S D P Y H A T T G P L D
15. R S D P Y H A T T G P L D
16. S D P Y H A T T G P L D
17. D P Y H A T T G P L D
18. P Y H A T T G P L D
19. Y H A T T G P L D
20. H A T T G P L D
21. A T T G P L D
22. T T G P L D
23. T G P L D
24. P L D
25. L D
26. D
27. .space

(10)

1. Q G P L G V K D R V K G R S D P Y H A T T G P L D
2. Q G P L G V K D R V K G R S D P Y H A T T G P L S
3. Q G P L G V K D R V K G R S D P Y H A T T G P L A
4. Q G P L G V K D R V K G R S D P Y H A T T G P L
5. Q G P L G V K D R V K G R S D P Y H A T T G P
6. Q G P L G V K D R V K G R S D P Y H A T T G
7. Q G P L G V K D R V K G R S D P Y H A T T
8. Q G P L G V K D R V K G R S D P Y H A T
9. Q G P L G V K D R V K G R S D P Y H A
10. Q G P L G V K D R V K G R S D P Y H

11. Q G P L G V K D R V K G R S D P Y
12. Q G P L G V K D R V K G R S D P
13. Q G P L G V K D R V K G R S D
14. Q G P L G V K D R V K G R S
15. Q G P L G V K D R V K G R
16. Q G P L G V K D R V K G
17. Q G P L G V K D R V K
18. Q G P L G V K D R V
19. Q G P L G V K D R
20. Q G P L G V K D
21. Q G P L G V K
22. Q G P L G V
23. Q G P L G
24. Q G P L
25. Q G P
26. Q G
27. Q
28. .space;

(11) D

1. P S K D C G D P K Y A Y F N G C S S P T A P L D P
2. P S K D C G S P K Y A Y F N G C S S P T A P L S P
3. P S K D C G A P K Y A Y F N G C S S P T A P L A P
4. S K D C G D P K Y A Y F N G C S S P T A P L D P
5. K D C G D P K Y A Y F N G C S S P T A P L D P
6. D C G D P K Y A Y F N G C S S P T A P L D P
7. C G D P K Y A Y F N G C S S P T A P L D P
8. G D P K Y A Y F N G C S S P T A P L D P
9. D P K Y A Y F N G C S S P T A P L D P
10. P K Y A Y F N G C S S P T A P L D P
11. K Y A Y F N G C S S P T A P L D P
12. Y A Y F N G C S S P T A P L D P
13. A Y F N G C S S P T A P L D P
14. Y F N G C S S P T A P L D P
15. F N G C S S P T A P L D P
16. N G C S S P T A P L D P
17. G C S S P T A P L D P
18. C S S P T A P L D P
19. S S P T A P L D P
20. S P T A P L D P
21. P T A P L D P
22. A P L D P
23. P L D P
24. L D P
25. D P
26. P
27. .space

(12) D

1. P S K D C G D P K Y A Y F N G C S S P T A P L D P
2. P S K D C G S P K Y A Y F N G C S S P T A P L S P
3. P S K D C G A P K Y A Y F N G C S S P T A P L A P
4. P S K D C G D P K Y A Y F N G C S S P T A P L D
5. P S K D C G D P K Y A Y F N G C S S P T A P L
6. P S K D C G D P K Y A Y F N G C S S P T A P
7. P S K D C G D P K Y A Y F N G C S S P T A
8. P S K D C G D P K Y A Y F N G C S S P T
9. P S K D C G D P K Y A Y F N G C S S P
10. P S K D C G D P K Y A Y F N G C S S
11. P S K D C G D P K Y A Y F N G C S
12. P S K D C G D P K Y A Y F N G C
13. P S K D C G D P K Y A Y F N G
14. P S K D C G D P K Y A Y F N
15. P S K D C G D P K Y A Y F
16. P S K D C G D P K Y A Y
17. P S K D C G D P K Y A
18. P S K D C G D P K Y
19. P S K D C G D P K
20. P S K D C G D P
21. P S K D C G D
22. P S K D C G
23. P S K D C

24. P S K D
25. P S K
26. P S
27. P
28. .space
1.

(13) D

1. D P S K D C G D P K Y A Y F N G C S S P T A P L D P M D
2. S P S K D C G S P K Y A Y F N G C S S P T A P L S P M S
3. A P S K D C G A P K Y A Y F N G C S S P T A P L A P M A
4. P S K D C G D P K Y A Y F N G C S S P T A P L D P M D
5. S K D C G D P K Y A Y F N G C S S P T A P L D P M D
6. K D C G D P K Y A Y F N G C S S P T A P L D P M D
7. D C G D P K Y A Y F N G C S S P T A P L D P M D
8. C G D P K Y A Y F N G C S S P T A P L D P M D
9. G D P K Y A Y F N G C S S P T A P L D P M D
10. D P K Y A Y F N G C S S P T A P L D P M D
11. P K Y A Y F N G C S S P T A P L D P M D
12. K Y A Y F N G C S S P T A P L D P M D
13. Y A Y F N G C S S P T A P L D P M D
14. A Y F N G C S S P T A P L D P M D
15. Y F N G C S S P T A P L D P M D
16. F N G C S S P T A P L D P M D
17. N G C S S P T A P L D P M D
18. G C S S P T A P L D P M D
19. C S S P T A P L D P M D
20. S S P T A P L D P M D
21. S P T A P L D P M D
22. P T A P L D P M D
23. T A P L D P M D
24. A P L D P M D
25. P L D P M D
26. L D P M D
27. D P M D
28. P M D
29. M D
30. D
31. .space

(14) D

1. D P S K D C G D P K Y A Y F N G C S S P T A P L D P M D
2. S P S K D C G S P K Y A Y F N G C S S P T A P L S P M S
3. A P S K D C G A P K Y A Y F N G C S S P T A P L A P M A
4. D P S K D C G D P K Y A Y F N G C S S P T A P L D P M
5. D P S K D C G D P K Y A Y F N G C S S P T A P L D P
6. D P S K D C G D P K Y A Y F N G C S S P T A P L D
7. D P S K D C G D P K Y A Y F N G C S S P T A P L
8. D P S K D C G D P K Y A Y F N G C S S P T A P
9. D P S K D C G D P K Y A Y F N G C S S P T A
10. D P S K D C G D P K Y A Y F N G C S S P T
11. D P S K D C G D P K Y A Y F N G C S S P
12. D P S K D C G D P K Y A Y F N G C S S
13. D P S K D C G D P K Y A Y F N G C S
14. D P S K D C G D P K Y A Y F N G C
15. D P S K D C G D P K Y A Y F N G
16. D P S K D C G D P K Y A Y F N
17. D P S K D C G D P K Y A Y F
18. D P S K D C G D P K Y A Y
19. D P S K D C G D P K Y A
20. D P S K D C G D P K Y
21. D P S K D C G D P K
22. D P S K D C G D P
23. D P S K D C G D
24. D P S K D C G
25. D P S K D C
26. D P S K D
27. D P S K
28. D P S
29. D P
30. D
31. .space

(13) pS

1. pSPSKDCGpSPKYAYFNGCSSPT
2. pSPSKDCGpSPKYAYFNGCSSP
3. pSPSKDCGpSPKYAYFNGCSS
4. pSPSKDCGpSPKYAYFNGCS
5. pSPSKDCGpSPKYAYFNGC
6. pSPSKDCGpSPKYAYFNG
7. pSPSKDCGpSPKYAYFNG
8. pSPSKDCGpSPKYAYFN
9. pSPSKDCGpSPKYAYF
10. pSPSKDCGpSPKYAY
11. pSPSKDCGpSPKYA
12. pSPSKDCGpSPKY
13. pSPSKDCGpSPK
14. pSPSKDCGpSP
15. pSPSKDCGpS
16. pSPSKDCG
17. pSPSKDCGpSPKYAYFNG
18. PSKDCGpSPKYAYFNG
19. SKDCGpSPKYAYFNG
20. KDCGpSPKYAYFNG
21. DCGpSPKYAYFNG
22. CGpSPKYAYFNG
23. GpSPKYAYFNG
24. pSPKYAYFNG
25. PKYAYFNG
26. KYAYFNG
27. AYFNG
28. YFNG

(14)

1. dSPSKDCGdSPKYAYFNGCSSPT
2. dSPSKDCGdSPKYAYFNG
3. FNG
4. .space;

(13) D

1. A P L D P M D P P G Y K L V T G D R N N S S C R N
2. A P L S P M S P P G Y K L V T G D R N N S S C R N
3. A P L A P M A P P G Y K L V T G D R N N S S C R N
4. P L D P M D P P G Y K L V T G D R N N S S C R N
5. P L D P M D P P G Y K L V T G D R N N S S C R N
6. D P M D P P G Y K L V T G D R N N S S C R N
7. P M D P P G Y K L V T G D R N N S S C R N
8. M D P P G Y K L V T G D R N N S S C R N
9. D P P G Y K L V T G D R N N S S C R N
10. P P G Y K L V T G D R N N S S C R N
11. P G Y K L V T G D R N N S S C R N
12. G Y K L V T G D R N N S S C R N
13. Y K L V T G D R N N S S C R N
14. K L V T G D R N N S S C R N
15. L V T G D R N N S S C R N
16. V T G D R N N S S C R N
17. T G D R N N S S C R N
18. G D R N N S S C R N
19. R N N S S C R N
20. N N S S C R N
21. N S S C R N
22. S S C R N
23. S C R N
24. C R N
25. R N
26. N
27. .space

(14) D

2. A P L D P M D P P G Y K L V T G D R N N S S C R N
3. A P L S P M S P P G Y K L V T G D R N N S S C R N
4. A P L A P M A P P G Y K L V T G D R N N S S C R N
5. A P L D P M D P P G Y K L V T G D R N N S S C R
6. A P L D P M D P P G Y K L V T G D R N N S S C
7. A P L D P M D P P G Y K L V T G D R N N S S

8. A P L D P M D P P G Y K L V T G D R N N S
9. A P L D P M D P P G Y K L V T G D R N N
10. A P L D P M D P P G Y K L V T G D R N
11. A P L D P M D P P G Y K L V T G D R
12. A P L D P M D P P G Y K L V T G D
13. A P L D P M D P P G Y K L V T G
14. A P L D P M D P P G Y K L V T
15. A P L D P M D P P G Y K L V
16. A P L D P M D P P G Y K L
17. A P L D P M D P P G Y K
18. A P L D P M D P P G Y
19. A P L D P M D P P G
20. A P L D P M D P P
21. A P L D P M D P
22. A P L D P M D
23. A P L D P M
24. A P L D P
25. A P L D
26. A P L
27. A P
28. A
